# Power-law behaviour of transcription factor dynamics at the single-molecule level implies a continuum affinity model

**DOI:** 10.1101/637355

**Authors:** David A. Garcia, Gregory Fettweis, Diego M. Presman, Ville Paakinaho, Christopher Jarzynski, Arpita Upadhyaya, Gordon L. Hager

## Abstract

Single-molecule tracking (SMT) allows the study of transcription factor (TF) dynamics in the nucleus, giving important information regarding the search and binding behaviour of these proteins in the nuclear environment. Dwell time distributions for most TFs have been described by SMT to follow bi-exponential behaviour. This is consistent with the existence of two discrete populations bound to chromatin *in vivo*, one non-specifically bound to chromatin (i.e. searching mode) and another specifically bound to target sites, as originally defined by decades of biochemical studies. However, alternative models have started to emerge, from multiple exponential components to power-law distributions. Here, we present an analytical pipeline with an unbiased model selection approach based on different statistical metrics to determine the model that best explains SMT data. We found that a broad spectrum of TFs (including glucocorticoid receptor, oestrogen receptor, FOXA1, CTCF) follow a power-law distribution, blurring the temporal line between non-specific and specific binding, and suggesting that productive binding may involve longer binding events than previously thought. We propose a continuum of affinities model to explain the experimental data, consistent with the movement of TFs through complex interactions with multiple nuclear domains as well as binding and searching on the chromatin template.

## INTRODUCTION

Transcription factors (TFs) are key regulatory proteins responsible for turning genes “on” and “off” by binding to enhancer or promoter elements across the genome (1). The current consensus describes TFs as being able to transition between three different states: 1) unbound from DNA (diffusing in the nucleus), 2) non-specifically bound and 3) specifically bound to chromatin (i.e. interacting with specific response elements) (2). However, biochemical studies and live-cell imaging experiments appear to disagree on the timescale that eukaryotic TFs can remain bound to chromatin, ranging from seconds to several hours or even days (3-8).

Advances in fluorescence microscopy have revolutionized our understanding of how TFs search and interact with chromatin (9). Single-molecule tracking (SMT), which is based on detecting and following through time the traces produced by the light emitted from a single fluorophore, allows the characterization of protein dynamics in live cells. When applied to the study of TFs, important information regarding the search and binding dynamics of these proteins can be extracted (9). SMT has been applied to over a dozen TFs, and has revealed that the time TFs remain bound to chromatin (i.e. residence time) is relatively short (seconds) and follows a bi-exponential distribution (reviewed in (2)). The bi-exponential behaviour is consistent with decades of biochemical studies, indicating that the DNA-bound population of molecules are composed of two distinct subpopulations: a short-lived fraction (‘fast stops’) and a longer-lived fraction (‘slow stops’). The fast fraction has been interpreted to represent non-specific binding to chromatin while the slow fraction is thought to be consistent with specific binding at enhancers or promoters (10-13). Experiments wherein TFs were mutated in their DNA-binding domains seem to confirm this model as the longer binding events were reported to be dramatically reduced (11,13-16).

However, this view is at odds with our current understanding of the nuclear environment. Far from being homogenous, the nucleus is highly compartmentalized and can impose constraints on the motion of many transcription-related molecules (7,17,18). For example, the presence of nuclear bodies, liquid-liquid condensates and distinct chromosomal architectures can critically affect the searching process of TFs for their target sites (6,19), implying that TF dynamics should exhibit dynamics beyond the bi-exponential model.

Recently, studies fitting TF dynamics to a three-exponential model have found longer residence times for the serum response factor (SRF) (over 4 minutes) (20) or CCCTC-binding factor (CTCF) (∼15 min) (21) than would be expected from a bi-exponential model. Moreover, a multi-exponential model was used to explain the dynamics of the TF CDX2 (22). Finally, the bacterial proteins, tetracycline repressor (TetR) and LacI, with no known endogenous targets in mammalian cells, show power-law behaviour when heterologously expressed (23,24). In fact, these non-specific binding events could be as long as specific ones (23).

A random variable *t* follows a power-law (25) for *t* > *t*_*min*_ if *f*(*t*) = *At*^−*β*^, where *A* is a constant and *β* is the exponent or scaling parameter. Power-laws are heavy tailed (right-skewed), which makes rare events more likely to occur than for exponential distributions; and *β* is a measure of the skewness. Many natural and artificial systems have been found follow power-law distributions (25). For proteins interacting with chromatin, it would mean that the frequency of binding events of a given TF will be inversely proportional to the residence time of said TF. In fact, binding times orders of magnitude longer than the average are likely to be observed. More importantly, for mammalian TFs that follow a power-law distribution, assigning discrete residence times for specific and non-specific binding would not be feasible. Whether this phenomenon occurs for endogenous mammalian TFs remains an open question. While these discordant results regarding TF binding dynamics could reflect the underlying biology, they may also arise due to the lack of consensus in the field regarding tracking algorithms, photobleaching correction methods, and model fitting.

Here, we revaluated some of the core aspects of the SMT technique, focusing on photobleaching correction methods. We then derived theory-based models for TF dynamics and a principled method to obtain optimal model parameters from empirical residence time distributions, using Bayesian statistics. We analysed the dynamics of several TFs, including the glucocorticoid receptor (GR), the oestrogen receptor (ER), the “pioneer factor” forkhead box A1 (FOXA1), the chromatin remodeler BRG1 (SMARCA4) as well as the architectural protein CTCF. Our data is consistent with power-law behaviour for all tested proteins. We further discuss theoretical considerations for how broad effective distributions of binding affinities can result in the observed power-law distribution. We suggest that TF dynamics is not explained by a simple separation between non-specific and specific binding but rather reflects the heterogeneous nature of chromatin structure and binding strengths.

## MATERIAL AND METHODS

### Plasmid constructs

The pHaloTag–GR has been previously described (13). The construct expresses rat GR fused with HaloTag protein (Promega, Madison, WI, USA) in the C-terminal domain under the CMVd1 promoter. The pHaloTag-H2B has also been previously described (26). The N-terminus of H2B is fused with the HaloTag. pHaloTag-H3 and-H4 were purchased from Promega (pFN21AE1298 and pFN21AE0273, respectively). The pHaloTag-ER and pHaloTag-FoxA1 has been previously described (27). The pHalo-CTCF expresses the mouse CTCF with HaloTag fused in the C-Terminal domain. It has been generated by PCR amplification from the pCTCF-GFP vector (28) and sub cloned into the pHalo-GR previously cut with PvuI and XhoI restriction enzymes (New England Biolabs). The pHalo-SMARCA4 was purchased from Promega (pFN21AE0798). The pSNAP and pSNAP-GR have been previously described (29).

### Cell culture and transfection

The 3617 mouse mammary adenocarcinoma cell line used in this study as well as the GR knock-out subclone expressing Halo-GR has been previously described (15,29). Cells were routinely cultured in high glucose DMEM supplemented with 10% fetal bovine serum and 2 mM L-glutamine at 37°C in a CO2-controlled humidified incubator. The cell line contains a stable integration of the rat GFP–GR under tetracycline regulation (30). To prevent expression of GFP–GR, the cells were grown in the presence of 5 µg/ml tetracycline (Sigma-Aldrich, St. Louis, MO, USA).

5 million cells were electroporated using BTX T820 Electro Square Porator (Harvard Apparatus, Holliston, MA, USA) in 100ul of DPBS with 2.5 ug of plasmid. 25 ms pulses of 120v were used and cells were resuspended in fresh media. Single-molecule imaging experiments were set up as follows: 100,000 electroporated cells were seeded onto each well of a 2-well Lab-Tek chamber (1.5 German borosilicate coverglass, Thermo Fisher, Waltham, MA, USA) in high glucose DMEM supplemented with 10% FBS (Life Technologies), 2mM L-glutamine, 5 µg/ml tetracycline, and cultured overnight. The media was then replaced with high glucose DMEM supplemented with 10% charcoal stripped FBS (Life Technologies), 2mM L-glutamine, 5 µg/ml tetracycline, and incubated at 37°C for at least 24 hours before labeling.

### Fluorescent labeling of Halo-tagged molecules and hormone treatments

Transfected cells were incubated with 5 nM JF549-HaloTag or 10 nM cpSNAP-tag (31) ligand for 20 min at 37°C. Stably integrated Halo-GR cells were incubated with 0.5 nM JF549-HaloTag for 20 min at 37°C. Free ligand was depleted by washing three times with phenol red free DMEM media (supplemented with 10% charcoal-stripped FBS and 5 µg/ml tetracycline) in 15 min intervals at 37°C. Next, cells were treated with 600 nM Corticosterone (Cort) (Sigma-Aldrich) or 100 nM Dexamethasone (Dex) (Sigma-Aldrich), or 100 nM oestradiol (Sigma-Aldrich) and incubated for 20 min at 37°C before imaging. For wash-out experiments, cells were washed with media three times for 4 different intervals (every 15 minutes for 1 hour or every hour for 4 hours) after 20 minutes of hormone treatment and finally imaged.

### Image acquisition for single-molecule tracking and analysis

A custom Highly Inclined and Laminated Optical sheet (HiLO) microscope was used as previously described in detail (29), with an objective heater to reduce drifting. Briefly, the custom-built microscope from the CCR, LRBGE Optical Microscopy Core facility is controlled by µManager software (Open Imaging, Inc., San Francisco, CA.), equipped with an Okolab stage top incubator for CO2 (5%) and temperature control (37°C), a 150X 1.45 numerical aperture objective (Olympus Scientific Solutions, Waltham, MA), a 561nm laser (iFLEX-Mustang, Excelitas Technologies Corp., Waltham, MA), and an acousto-optic tunable filter (AOTFnC-400.650, AA Optoelectronic, Orsay, France). Images were collected on an EM-CCD camera (Evolve 512, Photometrics). Tracking was performed in MATLAB (version 2016a, The MathWorks, Inc., Natick, MA) with the custom software TrackRecord [version 6, originally developed elsewhere (32) and updated in-house]. For step-by-step instructions, we will supply a **User Manual file** after peer-review. Briefly, in TrackRecord, to analyse each time series, data were filtered using top-hat, Wiener, and Gaussian filters, then particles were detected, fitted to two dimensional gaussian function for “super resolution” and finally tracked using a nearest neighbour algorithm (29). Particle trajectories are divided into mobile and immobile. The displacements of histones H2B characterize the thermal jiggling of the DNA and from it, two parameters are extracted called Rmin and Rmax. Rmin corresponds to the maximum displacement of 99% of histones at a time-lag of 2 frames (frame to frame displacement) while Rmax corresponds to the maximum displacement of 99% of histones at a time-lag of shortest track. The shortest track is calculated using the diffusion coefficient of GR (∼5 μm^2^/s) to minimize tracking errors as explained elsewhere (26). The immobile tracks are used to calculate the survival distribution using the Kaplan-Meier estimate. The 95% confidence interval was estimated using Greenwood’s Formula. All fits performed to the data were implemented with the nonlinear least square method using bisquare weights due to the noise on the tail of the survival distribution. Parameters used for acquisition conditions and analysis are shown in **Table 1**.

**Table 1.** Parameters used for each acquisition condition, and analysis of SMT data.

| Interval<br>(ms) | Exposure<br>(ms) | Laser<br>power<br>(mW) | Frame<br>Number | Maximum<br>(Pixels) | Shortest<br>track<br>(frames) | Gap<br>(frames) | Rmin<br>($\mu\text{m}$) | Rmax<br>($\mu\text{m}$) |
| --- | --- | --- | --- | --- | --- | --- | --- | --- |
| 200 | 10 | 0.96 | 600 | 4 | 4 | 2 | 0.23 | 0.31 |
| 200 | 100 | 0.29 | 900 | 4 | 4 | 2 | 0.21 | 0.29 |
| 500 | 500 | 0.16 | 800 | 4 | 2 | 2 | 0.23 | 0.29 |
| 1000 | 500 | 0.16 | 800 | 4 | 2 | 1 | 0.29 | 0.33 |

Exported tracking data was further analysed in MATLAB by a custom-made script (which will become available after peer-review). For comparison and control purposes, we also performed tracking using u-Tack (33). Briefly, we used the “Gaussian Mixture-Model Fitting” under default parameters for particle detection and localization. The tracking was then performed with the following values: Problem dimensionality = 2; Maximum Gap to close = 2; Minimum Length of Track Segment from the First Step = 4; Do segment merging = checked; Do segment splitting = checked. Finally, we chose the “Cost functions” and “Kalman Filter functions” to the “Brownian + Directed motion” model.

### Photobleaching correction

The modified correction method is based on histone data as a proxy for the fluorophore stability as originally performed elsewhere (16,34-36). One caveat still common to all methods described and applied here is the assumption of homogenous illumination, which unfortunately does not occur in HiLO set ups, as the laser hits the sample at an inclined angle [discuss elsewhere (29)]. A first step involves SMT of histones under the same conditions that the TF of interest will be imaged, as previously described (16,34-36). We tracked individual H2B, H3 or H4 molecules using HiLO by sub-optimal transient transfection of HaloTag-fused histones, labeled with JF549 HaloTag ligand. The three histone variants we tested presented statistically similar dynamics (**Figure S1A**). We continued with H2B for all further experiments. Histone genes are primarily transcribed upon entry into S-phase of the cell cycle (37). Due to our transient transfection approach, HaloTag-H2B proteins will be translated during interphase and therefore some histones will not be incorporated into chromatin at the time of acquisition (**Video S1**). Hence, the survival distribution of H2B will be composed of PB kinetics and a diffusive/transient binding component. To account for this behaviour and assuming PB kinetics at the single-molecule level is exponentially distributed, the survival distribution of H2B is fit to an exponential family with three components (**Figure S1B**). This constitutes the second step in the protocol, which only differ thus far from previous works in the fitting to three exponential rather than two-exponentials (16,34,35); or fitting to two exponential with an offset (36). The faster components characterize the dynamics of histones that have not been stably incorporated into chromatin, while the third (slower) component describes the PB kinetics of the fluorophore. The invariance of the first two components to photobleaching conditions strongly suggest they are indeed due to the dynamics of unincorporated histones, tracking errors and shortest track selection (**Figure S1C**). To confirm that the third component quantifies PB kinetics and not the intrinsic dynamics of H2B, we calculated PB lifetimes using histones H3 and H4 with the same statistical results (**Figure S1D**). Finally, the third step corrects the binding dynamics of the TF, that means, the ideal measurement where neither photobleaching nor sample drift occur, by using the experimental (observed) TF distribution and the PB dynamics. The “novelty” in our approach is that we use the third exponential distribution of H2B as a proxy for photobleaching, while other groups uses the entire H2B distribution (16). In this sense, no assumption on the survival distribution of the TF is made, and the empirical survival distribution is corrected by the third exponential component of the H2B survival distribution. More formally,

Let *P*(*τ*_*TF*_ ≥ *t*), *P*(*τ*_*p*_ ≥ *t*), *P*(*τ*_*TFreal*_ ≥ *t*) be the survival distribution of an experimental particle, photobleached particle and a dynamic particle, respectively. The survival distribution of an experimental particle is the one typically measured in a SMT experiment; the survival distribution of a dynamic particle corresponds to the ideal measurement where neither photobleaching nor sample drift occur. We are interested in *P*(*τ*_*TFreal*_ ≥ *t*)

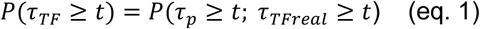

If a molecule is observed to live longer than *t* then it neither photobleached nor unbound from the DNA. These two processes are independent:

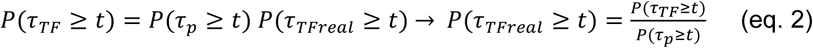

If the empirical survival distribution of photobleaching at the focal plane is available, then the dynamic survival distribution can be extracted from the microscopy data.

*P*(*τ*_*p*_ ≥ *t*) is estimated by fitting the survival distribution of H2B by a triple exponential function of the form:

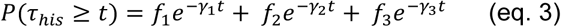

where *γ*_3_ corresponds to the photobleaching and *γ*_1_; *γ*_2_ the parameters of the dynamics of diffusive and/or unincorporated histones. The survival distributions are normalized with respect to the shortest track, for a shortest track of 6 frames and an acquisition interval of 200ms, the survival distribution is set up to *P*(*τ* ≥ 1.2 *s*) = 1.

Finally, assuming that the third component of *P*(*τ*_*his*_ ≥ *t*) corresponds to photobleaching:

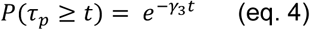

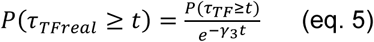

If we correct the H2B survival distribution with this method, we observe a predictable upward shift of the distribution (**Figure S1E**), in contrast to our previous methodology (15), wherein H2B data still artifactually resembles the dynamics of a TF. The high fluctuations at the tail of the distribution are likely due to noise in the data and the appearance of multiple particles within the point spread function, as illustrated in **Figure S1F**.

### Quantification and statistical analysis

For statistical analysis, all the parameters are reported by the ensemble average, standard deviation (s.d.) and number of observations. At least three biological replicates of SMT experiments were performed per condition. Two sample K-S test on the survival distribution were performed between replicates to confirm statistical reproducibility. Between 10 and 20 cells were imaged per SMT replicate. Each condition has at least 15000 tracks after analysis of SMT experiments. For survival distribution analysis, a statistical threshold of 5 tracks were implemented for visualization purposes only. Any point in the survival distribution with less than 5 cumulative tracks was not displayed in the figure. Data was not removed for fitting purposes. Fitting was done using non-linear least squares, initially a best local fit was found and then 50 iterations were run to find a global solution.

Simulations were written in MATLAB to numerically verify the different models of TF using the Gillespie algorithm (38). Graphical inspection was used to qualitatively determine if a straight line was observed for multiple decades in the case of a power-law fit in a log-log plot. Two different metrics were used to determine the difference between exponential models and power-law models. The first metric corresponds to Bayesian information criterion (BIC) using the probability distribution function (PDF) corrected for photobleaching as the likelihood function. BIC is a criterion for model selection that penalizes for model complexity (number of free parameters in the model). The PDF of TF dwell times was normalized between the minimum and maximum observation range (BIC1). BIC1 is given by (39):

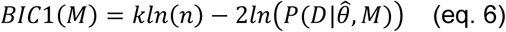

where *M* corresponds to the model (power-law, bi-exponential or triple-exponential), *k* corresponds to the number of parameters of the model, 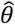 corresponds to the model parameters found by fitting, *D* the observed data and *n* the number of observations. 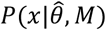 corresponds to the realization probability of *x* given the model PDF with parameters 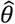. For SMT, *D* is the set of independent and identically distributed discrete experimental events and 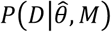 is calculated as follows:

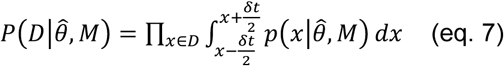

Where 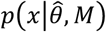 corresponds to the PDF of the model after photobleaching correction. For instance, the bi-exponential PDF is given by:

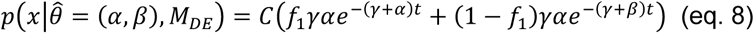

where *α, β* are the exponential parameters, *γ* the photobleaching rate and C a normalization constant.

The second metric, the evidence, in decibels (Db), for a particular model given the observed data and priors, was calculated to compare the alternative models explored. The evidence measures the probability of a particular model being the best predicting model in comparison with another model.

For instance, for the power-law model (*M*_*PL*_) the evidence versus the bi-exponential model (*M*_*DE*_) and the triple exponential model (*M*_*TE*_) is given by (40):

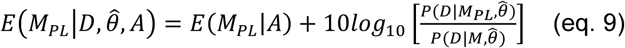

Where *M* = *M*_*DE*_ ∪ *M*_*TE*_

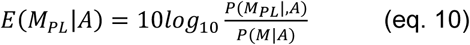

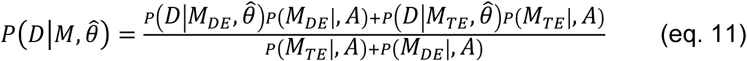

where *A* corresponds to the priors; *P, D* and 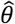 as defined for BIC1. Uniform priors were used for all model comparisons. For instance, an evidence of 30 Db corresponds to a probability higher than 0.999 that the power-law model better describes the data in comparison with an alternative model tested. In general, a positive value of the evidence indicates that the corresponding model is a better predictor of the data in comparison to the other tested models. If the evidence was not high enough to reach a conclusion about the comparison between the different models, more data was acquired until the evidence reached a satisfactory value.

Refer to **Table S1** for all statistical results. Table S1 lists the evidence in Db for the models, the difference of BIC1 (denoted as Delta-BIC1) between the power-law model and bi/triple exponential models. A positive value of the difference in BIC1 implies a preference for the power-law model over the bi- or triple-exponential models.

Model Selection was performed in the following manner: Graphical inspection for linearity of the survival distribution in Log-Log plot for at least 1.5 decades for power-law model consideration, the model with an evidence higher than 30Db and a difference of BIC1 in accordance with the model (a positive value for power-law as a better model, negative value for bi/triple exponential models) was chosen. Evidence does not take into account model complexity and therefore the model selection is done jointly with BIC1.

## RESULTS

### Photobleaching correction methods and their effect on survival distributions

When tracking TFs at the single-molecule level, the experimental information that is recovered is the time the molecule “remains” visible before it bleaches or moves out of the focal plane. Thus, binding events can be observed as stationary spots (**Figure 1A-C, Video S2A-D**). From these observations, one can obtain a local dwell time for TFs, which is defined as the time interval between a single molecule transitioning from a diffusive state to a bound state and its subsequent return to diffusion. The dwell time distribution is generated by integrating the ensemble-averaged distribution of bound times (**Figure 1D** and **Supplementary Note 1**.**1**). Most often, a “survival” distribution, defined as 1-CDF, where CDF is the empirical cumulative distribution function of dwell times, is used for further analysis [**Figure 1E**, GR dynamics adapted from (15)]. This plot represents the probability P that a molecule will last t number of time points, or longer. This survival distribution is fit to a bi-exponential distribution [**Figure 1E** and reviewed in (41)], and interpreted as the “three population model” (i.e. diffusive, fast bound or non-specific binding, slow bound or specific binding) as illustrated in **Figure 1F**. However, as can be seen in **Figure 1E**, the data shows a distinct departure from the bi-exponential fit, especially at longer dwell times.

**Figure 1.**
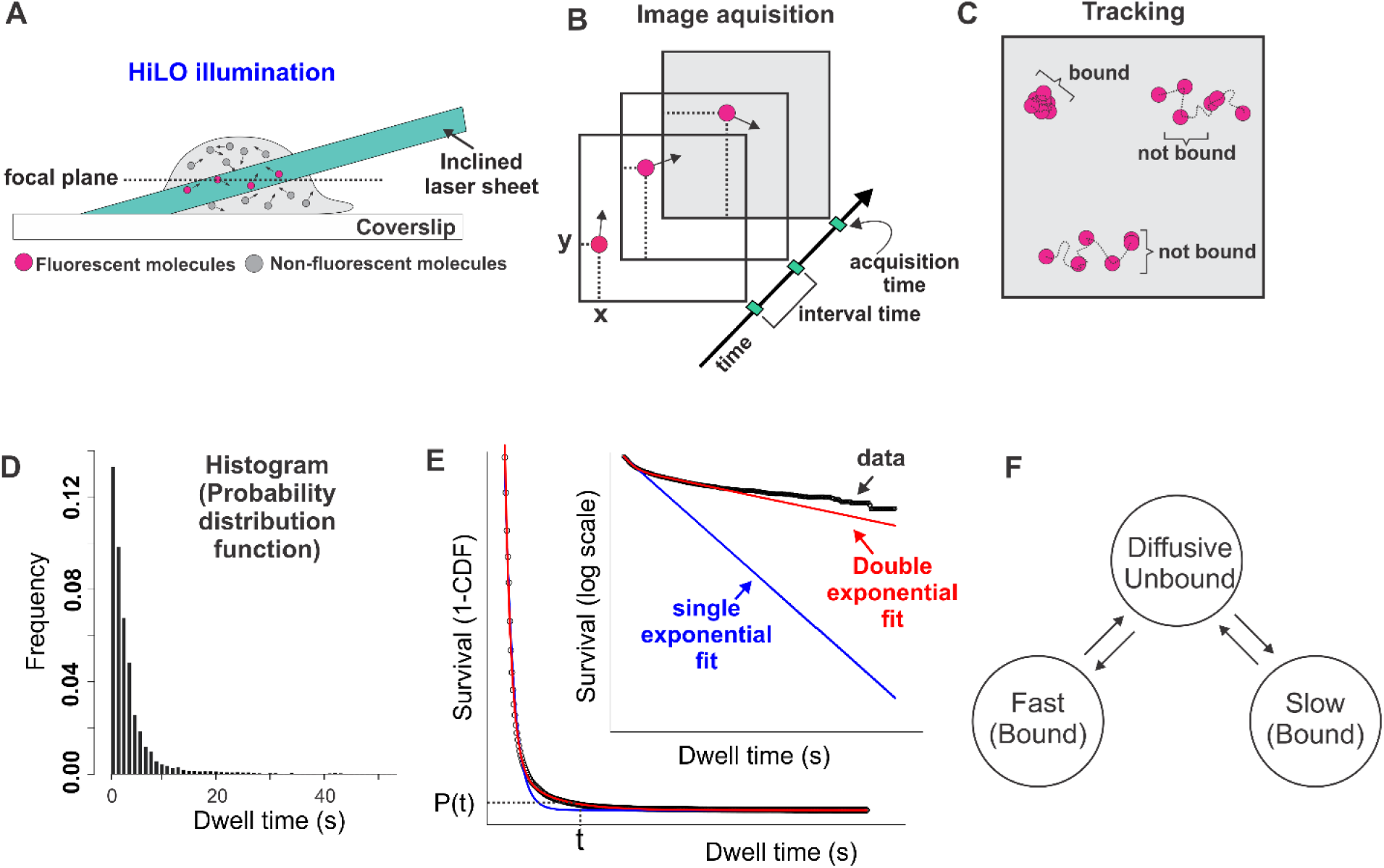
The current SMT pipeline and interpretation of TF dynamics. (**A**) A HiLO set-up is most commonly implemented to increase signal-to-noise ratio. A laser beam is tilted and hits the sample creating a thin illumination layer in the focal plane. (**B**) Several images are taken at specific yet variable acquisition and interval time conditions. (**C**) A tracking algorithm is used to follow individual molecules and classify them as either bound or unbound. (**D**) Histogram plotted from the bound population showing the frequency of TF molecules that are bound for a specific time (dwell time). Data acquired at 200ms interval for HaloTag-GR activated with its natural ligand corticosterone (15). (**E**) Fitting of the survival distribution (1-CDF; <u>c</u>umulative <u>d</u>istribution function) calculated from the data shown in D (circles) is fit to single-exponential (blue line) or bi-exponential (red line). Inset shows semi-log plot of the same. (**F**) Schematic showing the bi-exponential model according to which TFs occupy three different states: unbound from the DNA (diffusing in the nucleus), specifically bound (slow stops), and non-specifically bound (fast stops).

The upper temporal limit in SMT experiments is ultimately determined by the intrinsic photostability of the chosen fluorophore (42). When the affinity of bound TFs results in dwell times longer than those resulting from the average photostability of their fluorescent dyes, residence times cannot be resolved. Importantly, even when bound molecules have relatively lower affinities, they will appear to have shorter experimental dwell times due to photobleaching (PB) bias. To illustrate this known phenomenon, we conducted single-molecule imaging by transiently transfecting 3617 mouse mammary adenocarcinoma cells with the GR, a ligand-dependent transcription factor (43), tagged with HaloTag-Janelia Fluor 549 (JF549) (29) and stimulated with GR’s natural ligand corticosterone (Cort, 600 nM). When we artificially modulated the PB conditions by changing acquisition parameters (exposure time, imaging interval, laser power), the resulting kymographs have different typical lengths (**Figure S2 and Video S1**) and thus appear to have originated from different TFs. Therefore, PB must be properly corrected to prevent artifacts in the analysis of SMT data (32). Since PB correction methods vary widely among research groups (11,13,16,23,26,34-36,44) there is no standard approach to overcome the photobleaching bias of SMT strategies. Therefore, we decided to test the most common methods and our proposed approach by comparing how well they can correct the artifacts generated in GR dynamics measured with different acquisition conditions.

First, we tested the approach of estimating PB rates by counting, frame-by-frame, the number of particles of the TF of interest in the focal plane, then fitting the time-dependent decay of the molecule count (which is taken as a proxy for PB) to a bi-exponential model (10,13,15,26,45). This bi-exponential fit is finally used to normalize the survival distribution of the TF of interest, in this case, GR (**Figure 2A-B, method #1**). However, this method underestimates PB because most of the “counted molecules” are diffusive ones, and as such they are exposed to less laser illumination than bound molecules at the focal plane. Accordingly, this method fails to correct the apparent differences in GR survival distributions obtained from different acquisition conditions (**Figure 2B**).

**Figure 2.**
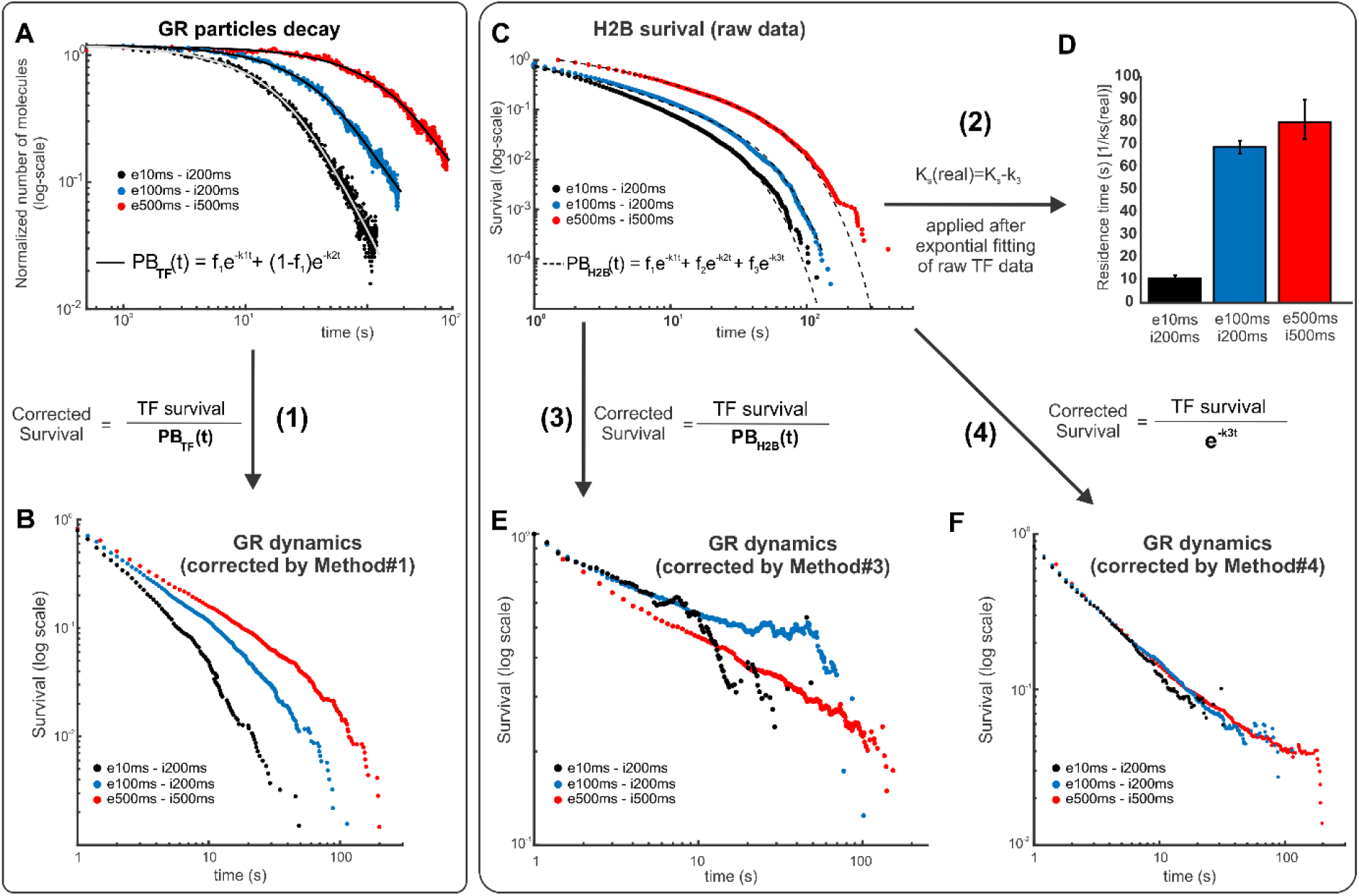
Effect of different photobleaching correction methods on the survival distribution of GR. (**A**) The number of particles (normalized to the initial number of particles for each cell at time zero) from frame-to-frame as a function of time shown for three different acquisition conditions as indicated in the legend (‘e’ denotes exposure time and ‘i’ denotes inter-frame interval). In method #1, this is taken as a proxy for photobleaching (PB), which is fitted to a bi-exponential function (black lines). (**B**) Effect of method #1 on GR dynamics at different acquisition conditions (e, exposure time; i, interval time). The corrected survival is obtained by dividing the observed TF survival to the bi-exponential distribution obtained in A. Number of cells/number of tracks are 67/9374 for GRe10ms/i200ms; 65/23172 for GRe100ms/i200ms; and 34/37953 for GRe500ms/i500ms. (**C**) In methods #2-4, the survival of H2B, taken under the same acquisition conditions as the TF, is used as a proxy for PB, which is fitted to either two or three family of exponentials. Number of cells/number of tracks are 100/36625 for H2Be10ms/i200ms; 63/40652 for H2Be100ms/i200ms; and 36/20307 for H2Be500ms/i500ms. (**D**) Method #2 does not correct the entire TF survival distribution but rather uses the slowest rate of the histone survival fitting (*k*_*3*_) to correct by subtraction the rate of the TF fitting (*k*_*s*_), thus obtaining the “real” rate (*k*_*s(real)*_). The panel shows the residence time (1/ *k*_*s*_) for the three experimental acquisition conditions. (**E**) Method #3 is similar to method #1, except that it uses H2B survival as a proxy for PB correction. The panel show GR dynamics at different acquisition conditions. (**F**) In Method #4, the exponential distribution of the slowest component in H2B survival is used as a proxy for PB correction. The panel show GR dynamics at different acquisition conditions. See **Table S1** for more data points details.

Another family of methods uses histones as a proxy for obtaining PB rates (**Figure 2C)**. Histones are a good representation of stably bound proteins because, after integration into chromatin, their residence time is much longer than the photostability of any currently available organic fluorophore (46). Therefore, by measuring the residence time of histones, one can obtain, in principle, a direct representation of PB for particles in the focal plane, since the disappearance of a long-lived particle will most likely represent a PB event. Different methodologies have been used under the umbrella of histone PB correction, ranging from measuring “bulk” histone levels and fitting the mean nuclear fluorescence (11), to variants of measuring histone dynamics at the single-molecule level (16,34-36). We will focus on the latter methods, as they use the same acquisition conditions as the TF of interest. One variant (34,35) fits the histone data to an exponential family (usually two components). However, instead of using the information of the entire histone survival distribution, only the decay rate of the longest component is used to correct the residence time of the TF by subtraction (**method #2**), effectively assuming that both TF and photobleaching dynamics follow exponential forms. Unfortunately, this method still gives different residence times for different acquisition conditions (**Figure 2D**). Another variant (16,36) is similar to method #1, but uses the survival distribution from histones instead of the number of molecules to normalize the TF data (**Figure 2E, method #3**). Although much better than method #1 (and #2), it fails to normalize GR distributions obtained with different acquisition conditions (**Figure 2E)** because the survival distribution of histones still has a significant population of diffusive molecules that are not incorporated into chromatin.

We therefore propose a modification to the previous PB correction methods, by combining the best of the three methodologies (**Figure 2F, method #4**, see methods for details). First, as in method #2 and #3, we fit the histone (HaloTag-H2B) SMT data, taken under the same imaging conditions as the TF of interest, to a family of exponentials. Second, we use the exponential distribution of the longer component (the entire exponential distribution rather than just the rate of the exponential) to normalize the TF survival data. In this way, we only correct for photobleaching by taking into account the bound histone population, without making any *a priori* assumptions about the survival distribution of a TF, as done in method #2. Using this modified version of PB correction, we find that GR survival time distributions obtained under different imaging conditions fall along the same curve as they should (**Figure 2F**). Taken together, our analysis suggests that this method more accurately corrects for photobleaching bias, as we obtain similar survival distribution of the TF irrespective of the photobleaching kinetics.

### GR dwell time distribution deviates from bi-exponential behaviour

We had previously used method #1 and characterized GR’s survival distribution as bi-exponential (15,27). Similarly, other groups have characterized other TFs as bi-exponentially distributed using their own PB correction methods [For example (7,11,14,34,36)]. Remarkably, when we apply our newly proposed method (**method #4**) of PB correction to the dwell time distribution derived from SMT data of HaloTag-GR activated with corticosterone (Cort, 600 nM), we find that the distribution now deviates from a bi-exponential distribution (**Figure 3A, Table S1**). The data look strikingly linear on a log-log plot (**Figure 3B**), which suggests power-law behaviour. The deviation from exponential is not due to an artifact of HaloTag, as dynamics of HaloTag-alone remain bi-exponentially distributed with no detectable “bound” molecules longer than 20 seconds (**Figure 3C**). To rule out artifacts from the imaging of exogenously expressed GR, we performed SMT on a GR knock-out cell line stably expressing Halo-GR at endogenous levels (15). The results are indistinguishable from the exogenous Halo-GR (**Figure 3D**), validating our transient expression strategy. Our data thus far suggests that a bi-exponential function does not describe GR dynamics at the single-molecule level, and a power-law might better explain the data.

**Figure 3.**
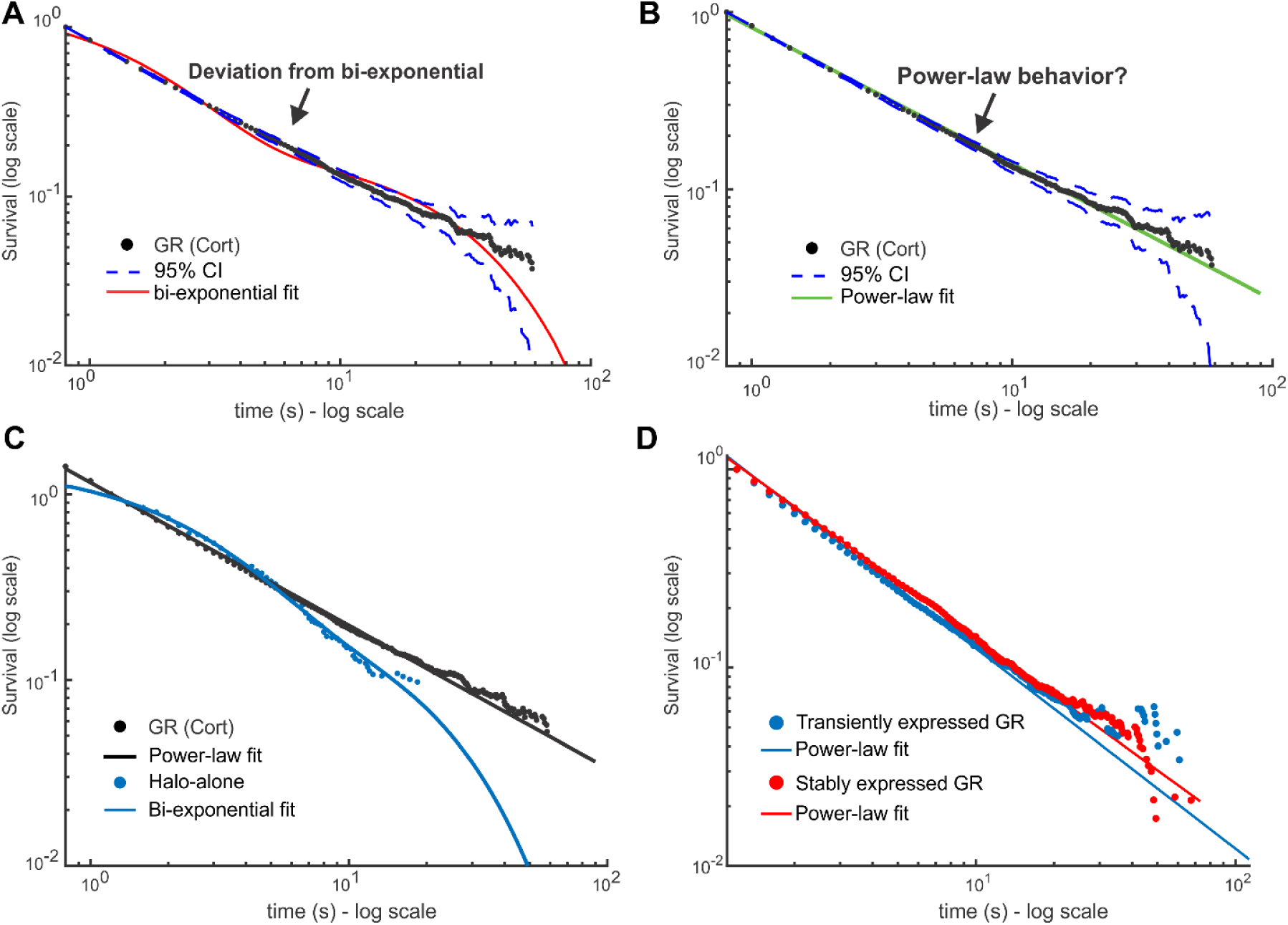
Impact of photobleaching correction on GR dynamics. (**A-B**) Single-molecule tracking data of GR activated with corticosterone (Cort). Data was acquired at 100 ms exposure time with 200ms interval. The survival distribution is shown (black), fit to a bi-exponential (A) or a power-law (B) function. Dashed lines show 95% confidence intervals (CI). Number of cells = 65; number of tracks = 23172. (**C**) Comparison of survival distributions of HaloTag-alone (blue) with a bi-exponential fit and HaloTag-GR (black) with a power-law fit. Data was acquired at 100 ms exposure time with 200ms interval. Number of cells = 64; number of tracks = 19436. (**D**) Survival distributions of HaloTag-GR, treated with Dex, either transiently transfected in 3617 cells (blue) or stably integrated in a GR knock-out subclone (red), expressed at endogenous levels. Data was acquired at 10ms exposure time with 200ms interval. Number of cells = 60; number of tracks = 7068 for GR transient. Number of cells = 60; number of tracks =16450 for GR stable. Coloured lines show power-law fits. See **Table S1** for details on fits.

### Theoretical models for TFs kinetics to interpret SMT data

Deviation from bi-exponential behaviour and the emergence of power-law behaviour has been described previously using heterologous expression of bacterial proteins (TetR and LacI) into mammalian cell lines (23,24). Moreover, a multi-exponential model has recently been proposed for the TF CDX2 (22) and SRF (20). To better understand the link between TF binding and the observed residence time distributions, we explored different theoretical models that may explain the emergence of different behaviours of the survival distribution.

Calculation of dwell time distributions is a first-passage time problem in stochastic analysis and has been widely used to characterize the kinetic properties of molecular motors and ion channels (47). When simple kinetic schemes are involved, dwell time distributions can be calculated analytically. However, for more complex systems, other methods must be used. One particularly powerful approach is to assign one or more states to “act” as an absorbing boundary, and then solve the associated first-order kinetic equations to obtain dwell time distributions (48) (**Supplementary Note 1**.**1**). We assume that the diffusive state (unbound) corresponds to an absorbing boundary state since tracked particles end with such transitions. The single molecule either photobleaches, disappears from the focal plane or begins diffusing. Any rebinding of the TF is considered an independent event.

We first examined the widely used bi-exponential model under this framework (**Figure 4A**). According to this model, TFs can occupy three different states: diffusive, slow and fast. In our analytic framework, the diffusive state plays the role of an absorbing boundary state, since particles entering the state are no longer tracked. The slow and fast states correspond to the empirically observed specific and nonspecific binding, respectively (reviewed in (41)). With this assumption of a well separated and narrow distribution of affinities, the expected behaviour of the survival distribution corresponds to a bi-exponential with the exponential parameters determining the average residence time of each state, as determined by analytic calculation (see **Supplementary Note 1**.**2)** and confirmed with stochastic simulations (**Figure 4B**) using the Gillespie algorithm (38). We note that this model does not allow for transitions between fast and slow states, which can be hard to interpret biologically, as searching (fast) should lead to specific binding (slow).

**Figure 4.**
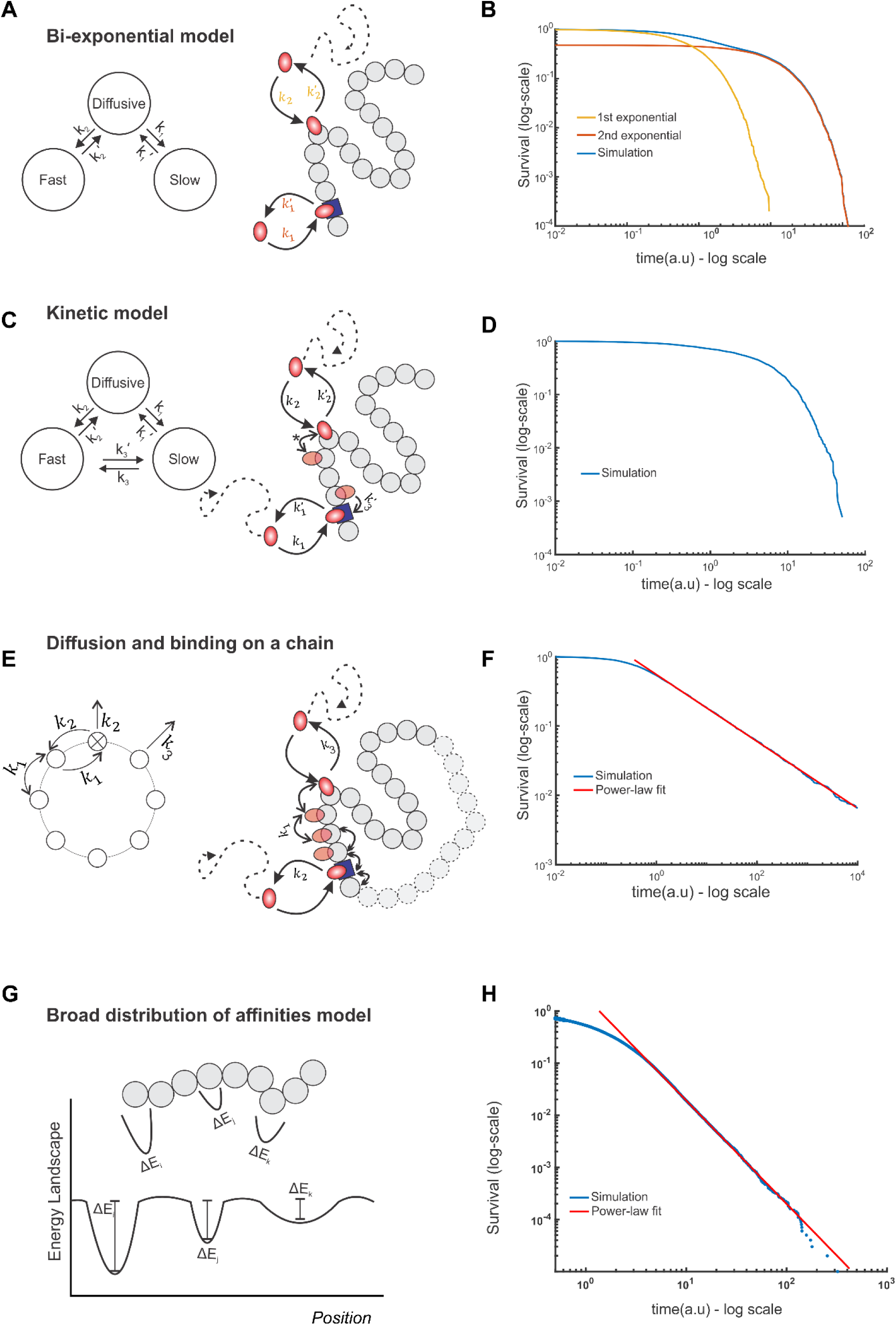
Theoretical models for TF kinetics. (**A**) State diagram (left) and schematic (right) of the bi-exponential model. TFs (orange oval) can bind to specific sites (blue square) or non-specific sites (grey circles) with rate constants *k*_*1*_ and *k*_*2*_ or unbind and return to the diffusive state with rate constants, *k*^*’*^_*1*_ and *k*^*’*^_*2*_ respectively (A). Transitions between specific and non-specific sites are forbidden. (**B**) Numerical simulation showing the emergence of bi-exponential behaviour for the model in A. The first and second exponential components are also shown as indicated. (**C**) State diagram (left) and schematic (right) of the Kinetic model. In addition to binding/unbinding to/from specific and non-specific sites, TFs can transition from specific sites to non-specific sites (with rate constant *k*_*3*_) and vice versa (with rate constant *k*^*’*^ _*3*_). Transitions between non-specific sites are considered indistinguishable (denoted by *). (**D**) Simulation results showing survival distributions arising from the kinetic model. (**E**) State diagram (left) of the continuum of affinities model, showing that transitions from a non-specific site to any other site occur with rate constant *k*_*1*_ and from a specific to a non-specific site with rate constant *k*_*2*_. Transitions to the diffusive state from the specific site occur with rate constant *k*_*2*_ and from a non-specific site with rate constant *k*_*3*_. Schematic (right) illustrating that a TF arrives at a random site and scans the DNA until it finds a specific site from which it can subsequently unbind. (**F**) Simulation of (E) to calculate the dwell time, which is defined as the time spent on the DNA, either bound or sliding, showing the emergence of power-law behaviour (red line, PL exponent 0.5, *k*_*1*_=10 a.u, *k*_*2*_=1 a.u, *k*_*3*_=10 a.u). (**G**) Schematic of the energy landscapes, representing the different binding affinities and the local microenvironment denoted as potential wells with different depths. (H) Numerical simulation of (G) showing the emergence of power-law behaviour (blue line). See also **Supplementary Note 1** for details.

We next extended the bi-exponential model to allow for transitions between the slow and fast components, which we call the kinetic model (**Figure 4C**). This model is a generalization of the bi-exponential model above. We note that due to the resolution limit (∼30nm), any transitions between specific and non-specific bound states cannot be distinguished experimentally. We found that for this extended model, the resulting survival distribution again corresponds to a bi-exponential distribution, with the exponential parameters as the eigenvalues of the transition matrix (**Supplementary Note 1**.**3**). Stochastic simulations were performed as before, and the resultant distribution, displayed in **Figure 4D**, again clearly demonstrates exponential behavior. An implication of the kinetic model is that simple interpretations of the exponential parameters as kinetic transition rates in either of the exponential models is not straightforward, since each rate constant might represent transitions between multiple hidden states and therefore the average dwell time may not necessarily represent the characteristic timescale of a particular interaction with chromatin.

Several theoretical studies have posited that TF searching for and “final” binding to its cognate site on the DNA involves a combination of bulk diffusion in the nucleus, 1D sliding along the DNA, hopping and translocation, and the theoretical search times for the TF to find specific sites in this framework have been estimated (49-51). In this model, TFs will have a multiplicity of short-lived bound states that must be accounted for in the analysis of dwell time data. To do so, we modelled TF movement on the DNA as hopping on a circular chain composed of specific and non-specific sites (**Figure 4E**). The main assumption in this model (**Supplementary Note 1**.**4**) is that the number of non-specific sites on the DNA is much larger than the number of specific sites. This is biologically reasonable as only a few to tens of thousands of specific sites are bound by any TF according to genome wide studies (52), while the entire genome contains millions of “other” potential chromatin sites. Since the length of time spent bound to the DNA depends on the number of non-specific sites visited before binding to and dissociating from the specific site, this will manifest itself as a continuum of effective binding affinities. An analytical solution can be found for the simplest case in which there is a single specific binding site and the TF can only unbind from this specific site (**Supplementary Note 1**.**4**.**3**). Biologically, this situation represents the case in which the TF finds the specific site and stays bound or rebinds rapidly upon dissociation. This has been hinted at by evidence of asymmetric diffusion prior to TFs binding (53) and IDR-IDR mediated phase separation of different transcription factors (54). A simulation based on the model gives rise to asymptotic power-law behaviour at time scales compatible with specific binding, for a number of representative parameter values (**Figure 4F**).

Finally, TFs can bind chromatin regions with varying physical microenvironments and motif degeneracy (55-57). These local properties affect the binding affinity of the TF. Given the heterogeneities in local organization and nuclear structure, TF binding sites on chromatin can be viewed as a collection of traps with a distribution of trap depths (**Fig. 4G**), analogous to binding affinities. If the binding affinities across different nuclear microenvironments and response elements are broadly and continuously distributed (for instance, exponentially distributed binding affinities), we can analytically demonstrate that the dwell times will asymptotically approach a power-law (58,59), as confirmed by simulations (**Fig. 4H** and **Supplementary Note 1**.**5**). In summary, we present phenomenological models that give us a framework to evaluate possible outcomes in SMT data.

### Dwell time distributions of GR and other TFs follow power-law behaviour

Having developed a theoretical framework to evaluate TF dynamic behaviour, we next explored which model better explains GR dynamics. We fit the survival distributions of GR activated with Corticosterone (GR-Cort, **Figure 5A**), or with dexamethasone (Dex, 100 nM), a more potent, synthetic hormone (GR-Dex, **Figure 5B**) to bi-exponential, kinetic and power-law models. As evident from the distributions, the bi-exponential and kinetic models show qualitative deviations from the data. We then used metrics based on the Bayesian information criterion (BIC) (60) test to choose the best predictive model (**see Methods**). Indeed, our statistical analysis confirms that a power-law corresponds to the best predictive model based on these metrics over a fit to the bi-exponential or kinetic model [Delta-BIC1 is 114423 (1047.3) for GR-cort; 13572 (942.8) for GR-Dex for the power-law fit compared to kinetic model (bi-exponential model)]. Moreover, the power-law fits were also superior to a tri-exponential fit (20,21) (see **Table S1** for all statistical comparisons). Surprisingly, we find that GR-Dex has a larger power-law exponent (β) than GR-Cort (*c*.*f*. **Figure 5A and 5B**), suggesting longer dwell times for the less potent ligand (Cort). This counterintuitive result is nevertheless consistent with a previous report correlating residence times with transcriptional bursting, wherein longer residence times (GR-Cort) correspond to a larger burst size, while overall transcriptional output is greater in GR-Dex due to a higher bound fraction (61).

**Figure 5.**
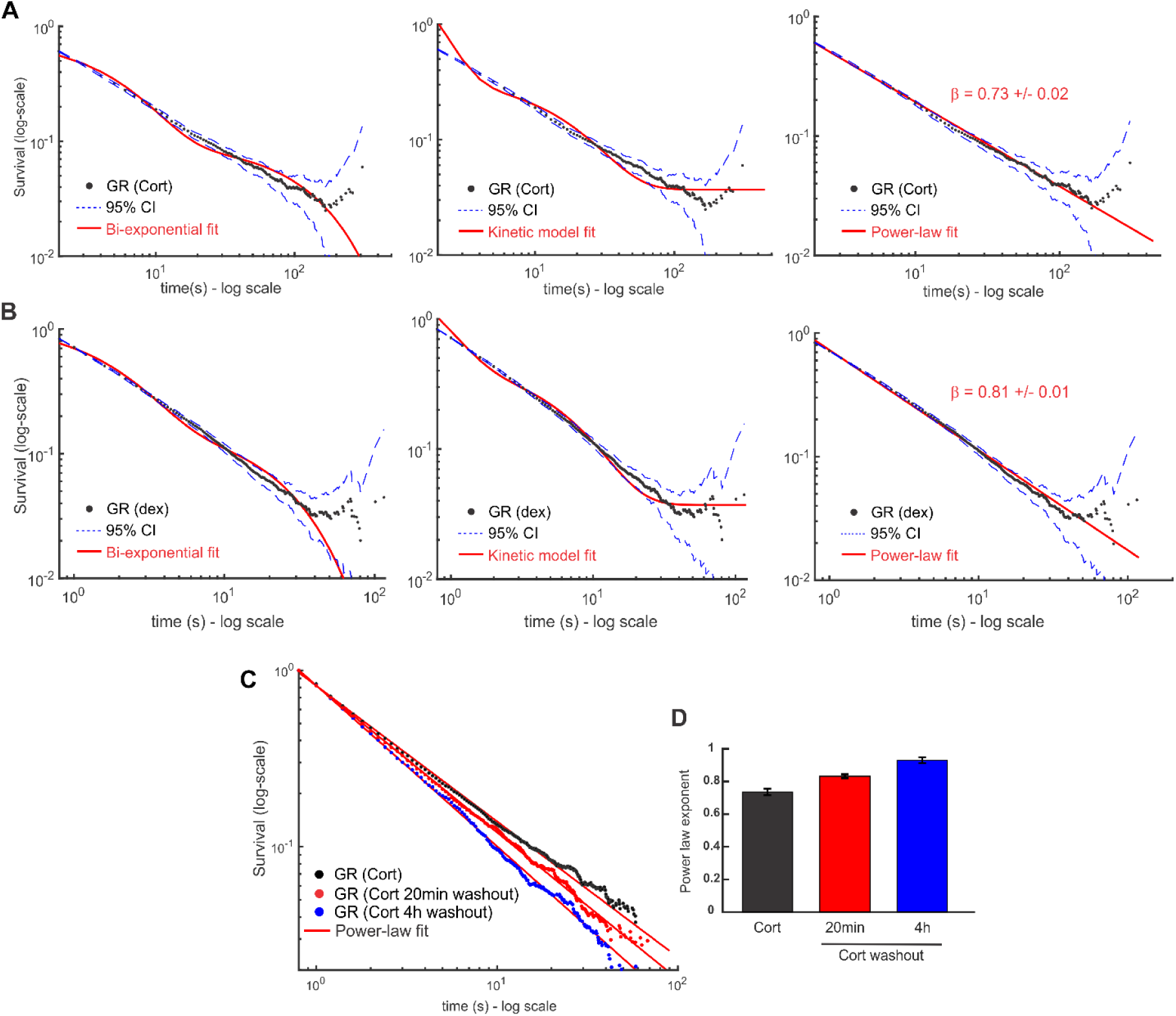
Dwell time distribution of GR follows power-law behaviour. (**A-B**) Survival distribution of GR activated with corticosterone (Cort, panel A, acquired with 500ms exposure time and 1000ms interval) and dexamethasone (Dex, panel B, acquired with 100ms exposure time and 200ms interval) obtained from SMT data. Number of cells/number of tracks are 30/15732 for GR (Cort); 40/29211 for GR (dex). Red lines show the best fit obtained for the bi-exponential model (left), the kinetic model (center) and a power-law (right). Dashed lines show the 95% confidence intervals (CI). (**C**) Survival distribution of GR activated by Cort (black symbols) or following washout of the hormone under a 20 min (red) or a more stringent 4h washout protocol (blue). Solid lines show fits to power-law model. Data acquired with 100ms exposure time and 200ms interval. Number of cells/number of tracks are 65/23172 for GR (Cort); 62/22530 for GR (Cort 20 min washout); 61/16611 for GR (Cort 4h washout). (**D**) Aggregate data for power-law exponents of fits to survival distribution of GR following stimulation by Cort, 20 min washout following stimulation and 4 h following washout. Errors represent 95% confidence interval. See **Table S1** for details on data acquisition and statistical comparisons.

We found that the power-law model better describes the data independent of the acquisition conditions (**Figure S3A**), yielding the same exponent under different PB rates (**Figure S3B)**. In contrast, a fit to a triple-exponential model showed that the parameters are dependent on acquisition times (**Figure S3C-D**). Further, the survival distributions obtained using a different tracking software [uTrack, (33)] were very similar to our tracking algorithm, ruling out any artifacts due to tracking (**Figure S3E)**. Finally, the power-law behaviour of the survival distribution of GR is conserved even if we use a different tag such as SNAP-Tag (62) (**Figure S3F**).

Previous work has largely assumed that the dynamics of non-specific binding is well described by a single exponential component with a much shorter dwell time than specific binding (11,13,26,29,34). However, heterologous proteins have also been reported to show power-law behaviour for the dwell times (23,24). To examine the dynamics of non-specific binding, we inactivated GR by washing out the hormone for 20 minutes, which greatly reduces specific binding as measured by chromatin immunoprecipitation (63). Interestingly, GR still exhibits power-law behaviour both for brief (20 min) washout, as well as longer washouts (four hours) (**Figure 5C**), although with shorter dwell times as indicated by a larger power-law exponent (**Figure 5D** and **Video S2E**).

To further establish the generality of our observations, we tested the dwell time distributions of different proteins previously characterized as bi-exponentially distributed by SMT (15,27,34). As with GR, both ER and FOXA1 exhibit power-law distributions (**Figure 6A**), with similar dynamics (β = 0.698 ± 0.005 for ER and 0.742 ± 0.003 for FOXA1) but slower compared to GR (β = 0.828 ± 0.004). This remains consistent with our previous observations wherein GR was more dynamic than ER and FOXA1 (27). Similarly, one of the major ATPase subunits of the SWI/SNF chromatin remodelling complex, SMARCA4, also exhibits a residence time distribution compatible with power-law behaviour (β = 0.845 ± 0.005, **Figure 6B**). Surprisingly, even the dynamics of the 11-zinc finger DNA-binding protein CTCF, involved in genome architecture among other functions (34), is better described by a power-law (β = 0.55 ± 0.02, **Figure 6B**). Taken together, our results indicate that the bi-exponential model might not properly reflect the dynamics of a wide range of chromatin interacting factors, and that it underestimates TF dwell times on chromatin. Thus, the power-law distribution emerges as a better descriptor of single-molecule dynamics, at least for the proteins tested.

**Figure 6.**
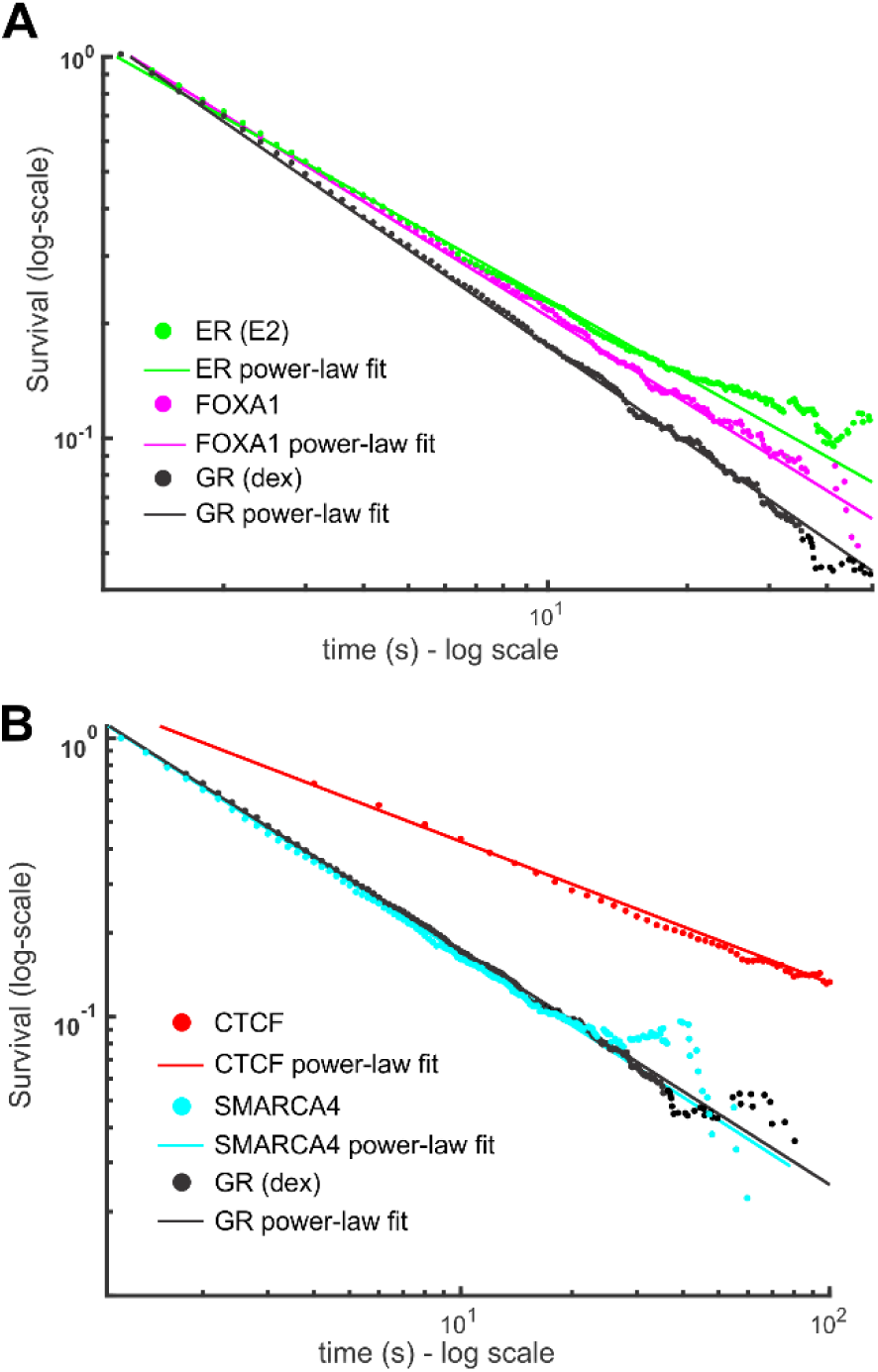
Dwell time distributions of TFs and other chromatin associated proteins show power-law behaviour. (**A-B**) Survival distribution calculated from SMT data of the Halo-Tagged oestrogen receptor (ER, activated with oestradiol, E2) (green), FOXA1 (magenta), CTCF (red) and SMARCA4 (cyan). GR (activated with dex, black) is shown for comparison in both plots. Data was acquired at 10ms exposure time with 200ms interval. Number of cells/number of tracks are 60/17823 for ER; 41/12864 for FOXA1; 50/7023 for SMARCA4, 40/29211 for GR (dex). CTCF data was acquired with a 10 ms exposure time and a 2000ms interval. Number of cells/number of tracks are 48/11606 for CTCF. Symbols are SMT data and solid lines are power-law fits to the data (see Table S1 for comparison and number of data points).

In conclusion, our analysis reveals hitherto unobserved features of the distribution of mammalian TF residence times (power-law vs. bi-exponential). This, in turn, suggests that specific and non-specific binding cannot be identified as two distinct populations with discrete (and measurable) residence times.

## DISCUSSION

In the present study, we propose a modified SMT pipeline with an improved photobleaching correction method to prevent bias of the dwell time distribution of TFs, and test underlying models using different statistical metrics. We are now able to reconcile data acquired under different experimental conditions whereas previous attempts were not successful (15).

We find that GR, as well as other TFs (ER and FOXA1), the chromatin remodeler SMARCA4 (also known as BRG1), and the insulator protein CTCF, all appear to exhibit power-law dynamics. It is generally accepted that to confirm this distribution, at least two orders of magnitude (both in x and y axes) should behave linearly on a log-log plot (64). This would require measuring TF binding up to several minutes (> 10 min), which is not currently feasible by SMT. Nevertheless, while there is a possibility that the power-law truncates at some point for really long binding times, we have enough statistical evidence to conclude that the power-law fit is a better predictor than a bi-exponential model over the observable experimental timescales. Examining whether more (or all) of the TFs originally characterized by bi-exponential behaviour are better described by a power-law exceeds the scope of this work and needs to be evaluated on a TF-by-TF basis.

Our observation of power-law behaviour of GR residence times suggests a model with a continuum of DNA-bound states rather than discrete non-specific/specific binding times (**Figure 7**). Consistent with this model, inactivation of GR by washing-out of the hormone revealed that the dwell time distribution also follows a power-law, indicating no apparent dynamical differences between specific and non-specific binding, as previously observed for bacterial proteins expressed in mammalian systems (23,24). Nevertheless, the overall residence times decrease when the receptor is less active, suggesting that a majority of the longer events observed with the fully activated receptor are associated with productive transcription as previously reported (11,13,15,16,26).

**Figure 7.**
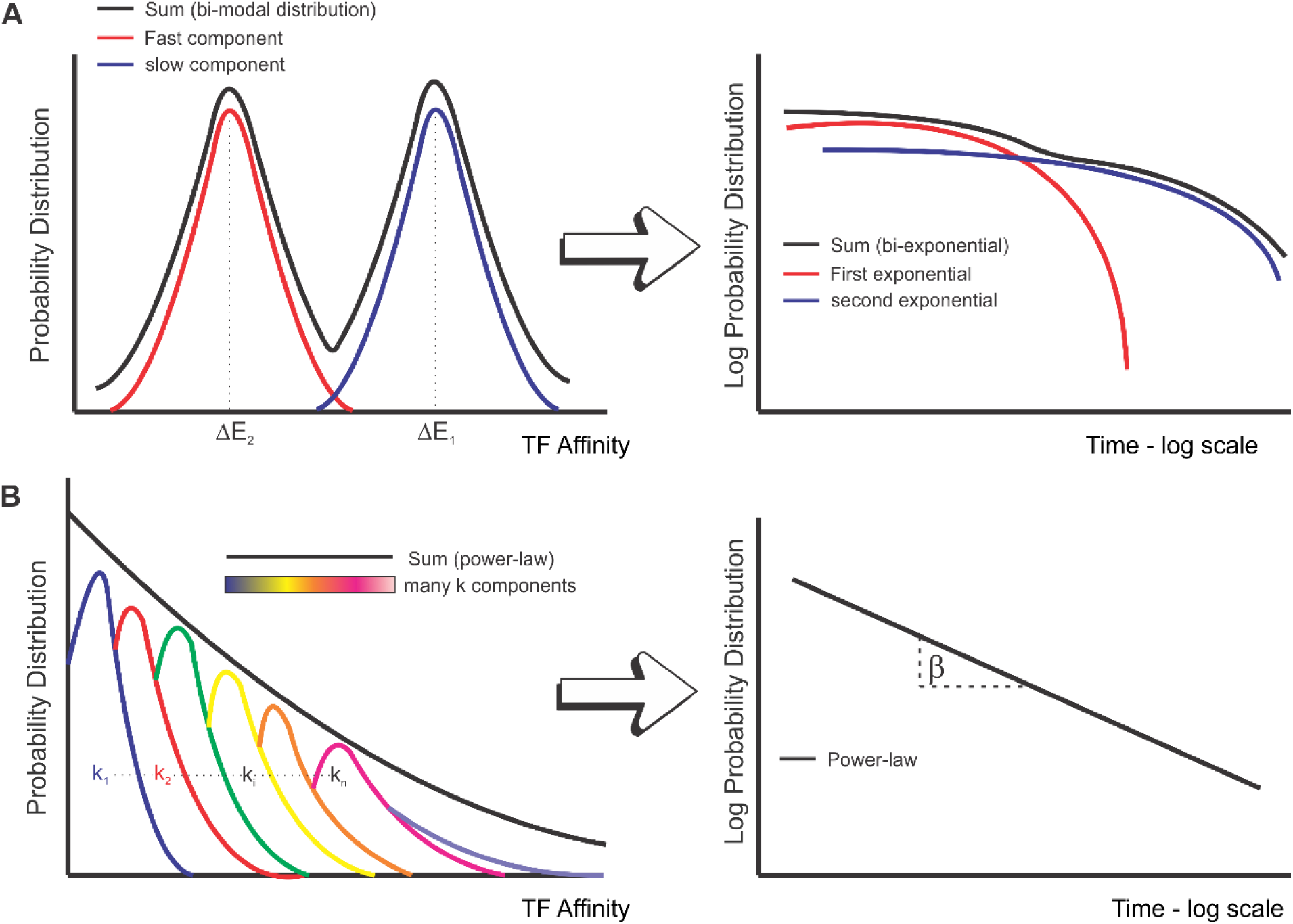
Heterogeneity in binding affinities can lead to a power-law behaviour of survival time distributions. (**A**) Schematic of the binding affinity distributions for a bi-exponential model. In this model, specific sites (blue) and non-specific sites (red) have a well separated and narrow distribution of affinities (ΔE, left graph), which results in a bi-exponential behaviour of the overall survival distribution (right graph, black curve). (**B**) Schematic showing a broad distribution of TF affinities (black line) which arises as a superposition of multiple sites with closely spaced affinity distributions (depicted in different colours in the left graph). Note that the distributions get progressively wider. This distribution of affinities may explain the emergence of power-law behaviour (characterized by the exponent, β) in the residence time of TFs (right graph).

An important characteristic of power-law distributions is that for exponents lower than or equal to one (as in our case), the mean is not a well-defined quantity (25). This implies that the mean can vary enormously from one measurement to the next and it is a limited measure of the process.

Interestingly, the heavy tails of power-law distributions imply that the probability of long-lived events is not negligible. This raises the possibility that productive binding events, although rare, may have dwell times much longer than previously appreciated, as indicated by the right-skewness of the distribution. We have recently shown a temporal correlation between GR dwell times and bound fraction with the length and frequency of transcriptional bursting (61). A similar behaviour has been observed in yeast with the Gal4/GAL3 model (65). However, non-specific binding can also result in TF binding events with long residence times, the implications of which are still not known. Critical efforts are required to investigate whether the slow(er) stops seen in SMT are matched exclusively to specific interactions with chromatin (7). For example, GR binding precedes RNA synthesis by ∼3 min (61). Alternatively, a sub-population of these “stops” could correspond to microscopic regions in the nucleus where diffusion is severely limited, due to transient interaction with “clustered” structures such as foci observed for GR (66), or another hitherto unknown mechanism.

The emergence of power-law might reflect the wide distribution of binding affinities in the nucleus. This broad distribution of affinities is puzzling but may be explained by a diverse set of non-mutually exclusive mechanisms. First, nuclear structure and the chromatin environment is known to be highly heterogeneous (9,17). Thus, TFs will encounter a wide variety of chromatin states (compacted fibers, different nucleosome modification conditions, etc.). Moreover, affinities for the thousands of alternative binding sites in response elements likely vary significantly. Furthermore, recent work points to the presence of transcriptional hubs and liquid-liquid phase separation domains (54,67-70) that contribute to the complexity of nuclear organization. If TFs exhibit different dynamical properties in these structures, it would not be surprising to find a broad variation in binding affinities. Second, TFs with intrinsically disordered regions (IDR) could adopt a broad distribution of conformations due to the diversity of protein folding (71), hence accounting for a broad distribution of binding affinities (55). An interesting corollary of this model is that TF lacking IDRs should not exhibit power-law behaviour if binding heterogeneity is coming from protein-protein interactions. Third, power-law distributed dwell time distributions can emerge as a consequence of the molecular kinetics of the protein itself, as recently reported *in vitro* by RNA polymerase II in bacteria (72). Fourth, the heterogeneity in the searching mechanism of TFs may affect the effective affinity constant observed in SMT experiments. In support of the latter, while heterologous expression of TetR in mammalian cells showed power-law behaviour for non-specific binding, it could still be described as an exponential on an artificially (and single) specific DNA binding array (23). Thus, the intrinsic nature of the searching mechanism of any DNA-binding protein in native chromatin may be governed by power-law dynamics. In addition, the heterogeneity of dwell times in the thousands of response elements for an endogenous TF could explain why GR can exhibit power-law tails as opposed to TetR, which can only bind to one artificial array site. Interestingly, a study in yeast (73) reports that both the TF Ace1p and the chromatin remodeler RSC binding follow a bi-exponential binding distribution in cells containing a natural tandem of ten CUP1 (Ace1p responsive) genes. This dynamic and discrete behaviour, in contrast with our GR data, can be explained by the particular and homogeneous chromatin environment of a single array of specific sites. Consequently, we speculate that a broad distribution of binding affinities due to a whole population of different binding sites (thousands in the case of GR) may result in power-law behaviour (**Figure 7**). Overall, independent of the mechanism behind power-law behaviour, the resulting broad distribution of binding affinities in these scenarios goes against the widely held assumption that TF dynamics on chromatin results from well-separated and narrow distributions of specific and non-specific binding with well-defined binding times (**Figure 7**).

While SMT methodology gives us the opportunity to study TF dynamics with unprecedented temporal and spatial resolution, it still has some major drawbacks. The sparse labelling conditions needed to resolve individual molecules severely limits the possibility of following all functional TFs at a time, and therefore may affect the implementation of a two-colour version where two different proteins interact at the single-molecule level. In addition, we still do not have direct measurements of the affinity at specific sites which makes it difficult to functionally distinguish between specific and non-specific binding. Nevertheless, the current major limitation in SMT is the photostability of the fluorophore, which limits the dynamical range of experiments and prevents accurate analysis of long TF trajectories that sample over different binding and/or diffusive events. Our temporal measurement window will improve with better, more stable fluorophores. Until then, our proposed pipeline allows us to have better estimates on the dynamics and the residence time distribution of TFs.

In summary, by incorporating an improved PB correction method and testing different models, we showed that the survival distribution of GR and other TFs dwell times does not follow an exponential model. Ultimately, if there is a way to define or distinguish non-specific from specific binding, our results indicate that it cannot be based on their global residence times. However, the data is consistent with a power-law distribution, which we suggest may arise generically due to heterogeneities in TF interactions with DNA or in the diffusive environment in the nucleus. Thus, the slope of the residence time distribution does provide an estimate of the overall affinity and can be used to compare TFs and their function under different conditions.

## AVAILABILITY

Tracking was performed in MATLAB (version 2016a) with the custom software TrackRecord (version 6). The latest version of this software, together with a few examples’ datasets, will be freely available at GitHub after peer-review. Exported tracking data was further analysed in MATLAB by a custom-made script which will be also available at GitHub.

## ACKNOWLEDGEMENT

We thank Tatiana Karpova and David Ball from the Optical Microscopy Core at the NCI, NIH for the assistance in imaging and data processing. We thank Luke Lavis (Janelia Research Campus) for providing HALO dyes. We thank Kaustubh Wagh for helpful discussions of the analytical results and Anders Sejr Hansen for his useful comments on the first version of the pre-print.

## FUNDING

This work was supported (in part) by the Intramural Research Program of the National Institutes of Health, National Cancer Institute, Center for Cancer Research. D.M.P was supported, in part, by the National Scientific and Technical Research Council (CONICET). V.P. was supported, in part, by the Academy of Finland, the University of Eastern Finland strategic funding and the Sigrid Jusélius Foundation. A.U. acknowledges support from the National Cancer Institute-University of Maryland (NCI-UMD) Cancer Technology Partnership, and the awards NSF PHY 1607645, NSF PHY 1806903, NSF PHY 1915534.

## CONFLICT OF INTEREST

The authors declare no competing interests

## Notes

### Competing Interest Statement

The authors have declared no competing interest.

