## Supplemental Figures, legends and Notes for "Power-law behaviour of transcription factor dynamics at the single-molecule level implies a continuum affinity model"

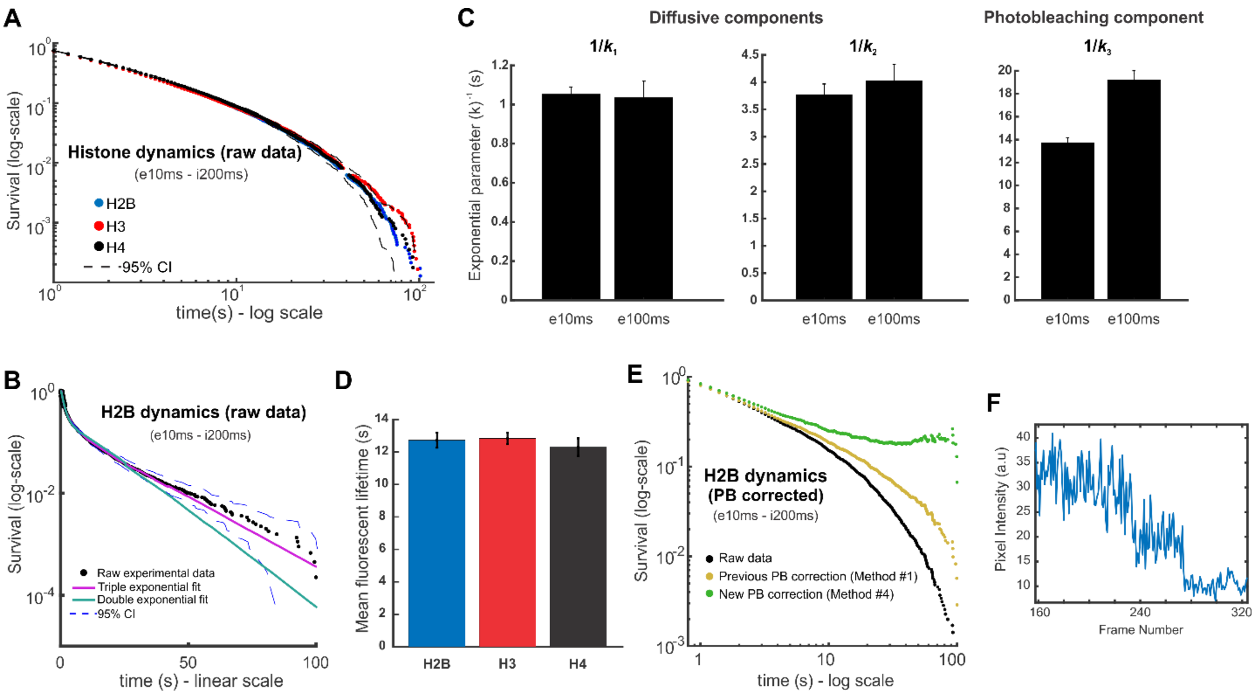


**Figure S1**. Histone dynamics as a proxy for photobleaching correction. (**A**) Survival distribution of histones H2B, H3 and H4 acquired under the same acquisition parameters as indicated (e, exposure time, i, interval). Number of cells/number of tracks are 100/36625 for H2B; 59/11708 for H3; 43/15601 for H4. (**B**) Fit of H2B survival distribution to a double exponential and triple exponential. A triple exponential better represents the experimental data where the slower component corresponds to the photobleaching rate in the focal plane. CI is the confidence interval. (**C**) Fitting the H2B data from two different exposure conditions (10ms and 100ms) to a triple exponential model gives the exponents $k_1$, $k_2$, $k_3$. The bar graph shows the mean +/- 95% confidence interval. (**D**) Mean fluorescence lifetime calculated as 1/ $k_3$ where $k_3$ is the slowest rate of the triple exponential. H2B (12.73 +/- 0.46 s), H3 (12.84 +/- 0.35 s) and H4 (12.30 +/- 0.55 s). Errors represent 95% confidence interval. (**E**) Survival distribution of H2B dynamics (black) corrected with our previous correction method (yellow), or the upgraded one (green). Note that after correction with method #1, H2B still has a finite dwell time. However, after correction with method #4, H2B presents two different regimes: stably incorporated histones that have very long residence times (plateau) and a dynamic regime representing unincorporated histones nonspecifically interacting in the nucleus. (**F**) Representative intensity profile of a histone particle selected from the tail of the distribution, with a cumulative probability of less than 1%. These very long events (outliers) usually present multiple photobleaching steps, indicating multiple particles at the same point-spread function. This might explain the deviations in the long tail of the H2B distribution after photobleaching correction. See **Table S1** for data points details

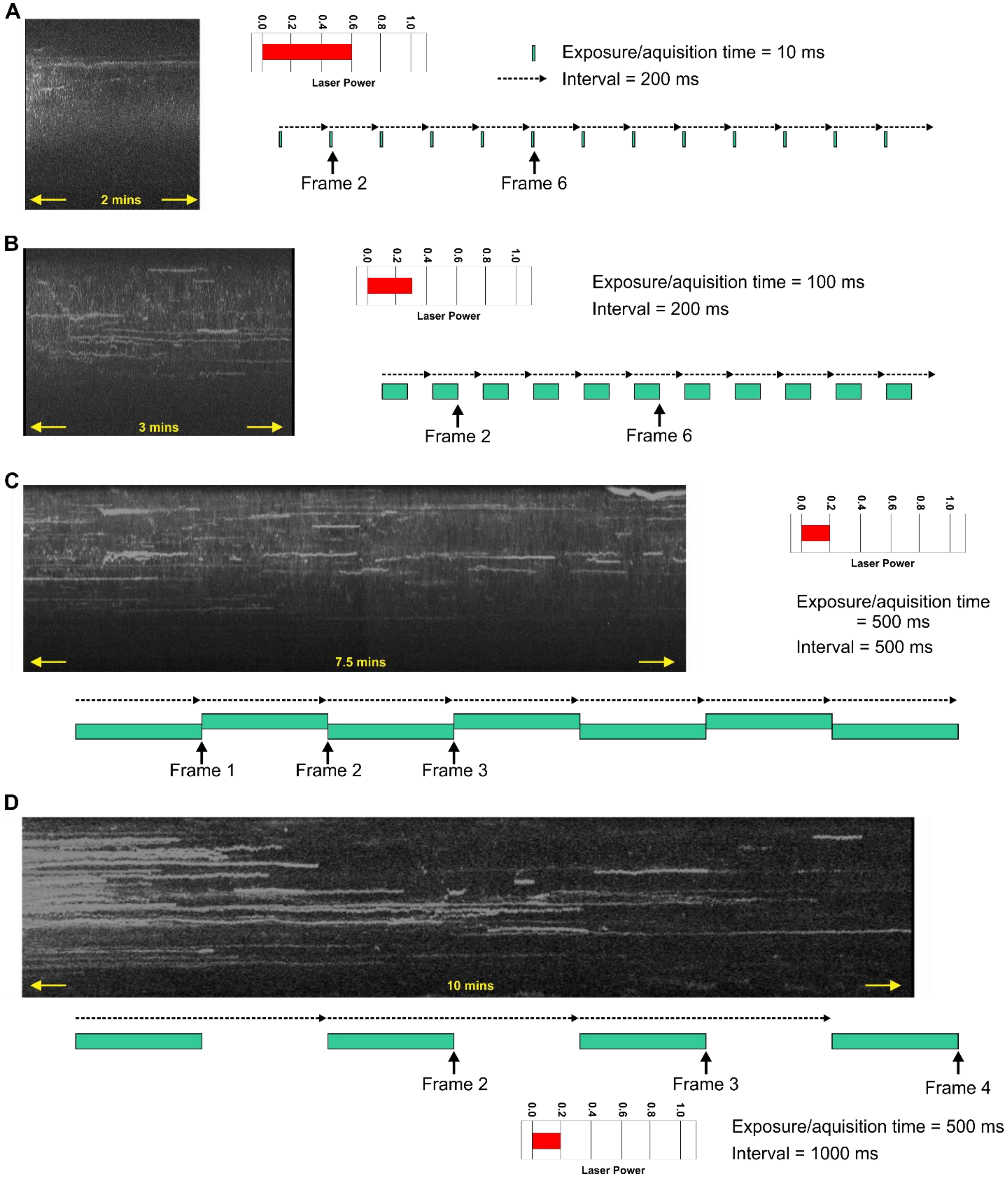


**Figure S2**. Qualitative illustration of photobleaching artifacts in SMT experiments. Single-molecule tracking data of the glucocorticoid receptor (GR) activated with corticosterone (Cort). The figure shows representative kymographs of GR molecules taken at different acquisition conditions (**A-D**), as indicated. The figure shows that track lengths are dependent on photobleaching kinetics, artificially modifying the apparent dwell time of GR. Without further analysis, the kymographs resemble different TFs.

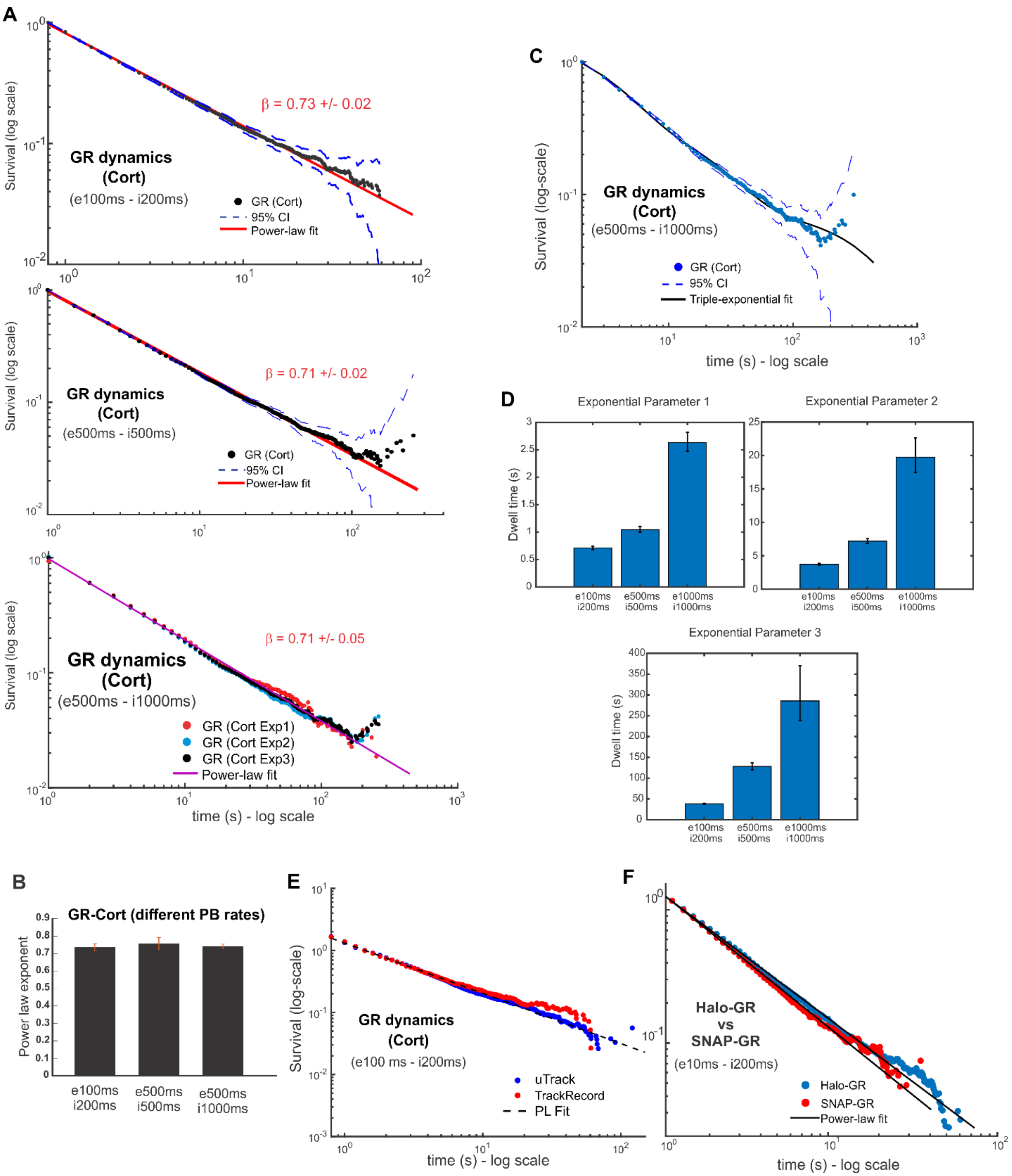


**Figure S3**. GR dwell times follow power-law distribution. (**A**) Power-law fits (red line) to the survival distribution of Corticosterone (Cort) activated GR (black symbols) from SMT data acquired at the indicated exposure (e) and interval times (i). Number of cells/number of tracks are 65/23172 for GRe100ms/i200ms; 34/37953 for GRe500ms/i500ms; and 30/15732 for GRe500ms/i1000ms. Bottom panel shows independent replicates of Cort-treated GR data acquired at 1000ms intervals, exemplifying reproducibility between SMT experiments. (**B**) Power-law exponent of GR-Cort SMT data under different photobleaching rates, generated by modulation of acquisition conditions, as indicated. Error bars represent 95% confidence intervals. (**C**) Survival distribution of GR-Cort fit to a triple exponential function. (**D**) The three exponential parameters of the triple exponential fit for different acquisition conditions as indicated shows that these parameters depend on acquisition conditions. (**E**) Survival distribution of GR dynamics tracked with uTrack (bue symbols) or TrackRecord (red symbols) software packages. Dashed line shows the power-law fit for uTrack. Number of cells/number of tracks are 65/23172 for TrackRecord; 40/11890 for uTrack. (**F**) Survival distribution of GR either tagged with HaloTag (Halo-GR, blue symbols) or SnapTag (SNAP-GR, red symbols) and the corresponding power law fits (black line). Number of cells/number of tracks are 67/9374 for Halo-GR; 50/7023 for SNAP-GR. See **Table S1** for details on data points and statistics.

**Table S1**. Data acquisition and statistical results. The first tab (TF_Fitting) shows for the indicated TF: the acquisition interval (in ms), exposure time (in ms), number of tracks, the evidence for the models (in Db), and the difference of BIC1 (denoted as Delta-BIC1) between the power-law model and bi/triple exponential models. Since the kinetic model is a special case of the bi-exponential model, model comparison was limited to the power-law, bi-exponential and triple-exponential models. The second tab (Histone_Fitting) shows the same information for H2B, H3, and H4 data. For histones, model comparisons were done between the bi-exponential and triple exponential models only.

**Video S1.** Single-molecule tracking data of HaloTag-H2B. Representative time-lapse movies of H2B in cells activated with Cort (A,B,C,D).The acquisition conditions are displayed above each movie. The movies are played 5X real time.

**Video S2**. Single-molecule tracking data of HaloTag-GR. Representative time-lapse movies of GR activated with Cort (A,B,C,D), or activated with Cort then washed-out for 4 hours (E). The acquisition conditions are displayed above each movie. The movies are played 5X real time.

### Supplementary Note for "Power-law behaviour of transcription factor dynamics at the single-molecule level implies a continuum affinity model"

#### 1 Theoretical models for TF survival distribution

In an SMT experiment, the protein of interest is tagged with a fluorescent probe and imaged. Binding events are then associated with stationary particles in the focal plane. The final experimental information that can be recovered is the time that a protein can be detected in the imaging volume before it bleaches, or moves out of the focal plane. From these observations, one can calculate a dwell time for transcription factors (TFs) which is defined as the time interval between a single molecule transitioning from a diffusive state to a bound state and its subsequent unbinding from DNA and return to the diffusive state. The dwell time distribution is obtained by calculating the ensemble distribution of bound times for a specific TF in different cells in the experiment after photobleaching correction (see Methods).The survival distribution is then calculated as 1-CDF, where CDF is the empirical cumulative distribution function of dwell times.

##### 1.1 Absorbing Boundary State Method

Calculation of dwell time distributions is a first-passage time problem in stochastic analysis and these distributions have been widely used to characterize kinetic properties of molecular motors and ion channels (1). In cases involving simple kinetic schemes, the dwell time distributions can be calculated analytically but for more complex schemes, a number of methods have been utilized. One particularly powerful approach is to assign one or more states to act as an absorbing boundary and then solve the associated first-order kinetic equations to obtain dwell time distributions. We assume that the diffusive (unbound) state corresponds to an absorbing boundary state, since the measurement ends with such transitions, because the particle either photobleaches, disappears from the focal plane or begins diffusing; any rebinding of the TF is considered as an independent event. This assumption implies that the population of particles in the absorbing boundary state increases with time. At the end of every experimental measurement, all the observed TFs transition to the absorbing state since the experiments are continued until most particles are bleached.

For a general process, a TF can be found in any state $i$ such as diffusing around the nucleus, confined in a microenvironment, bound to a particular specific or non specific site of the DNA. When a TF transitions to a diffusive state, it cannot be observed experimentally and this state plays the role of an absorbing boundary state. We observe the system over a time interval from $t' = 0$ to $t' = \tau$, during which individual TFs may undergo transitions between different states $i \in \{1, ..., n\}$. When a transcription factor is in a “bound” state, it can be observed experimentally as a trace as displayed in Supplementary Note Fig. 1a. Each TF in any of these bound states will be experimentally recorded from a certain time interval $t_1$ to $t_2$ with $t_2 - t_1 < \tau$. $t_1$ corresponds to the time when the transcription factor transitions to a “bound” state and $t_2$ corresponds to the time when the TF enters an absorbing state (diffusion in the nucleus). All the traces will be shifted by a time $t_1$ to a new aligned time $t = t' - t_1$ so that all the TFs begin in a bound state at $t = 0$ (Supplementary Note Fig. 1b). During the experimental time $\tau$, a finite number of TFs ($N$, equal to the number of traces) will be observed. When a TF transitions to a diffusive state, it cannot longer be observed experimentally and this state plays the role of an absorbing boundary state.

Let $p_i(t)$ correspond to the probability of being in state $i$ at time $t$.$\sum_{i \in boundaries} p_i(t)$ corresponds to the population of all absorbing states. To calculate number of unbinding events over a certain time interval ($f(t)$ dwell-time distribution, Supplementary Note Fig. 1c - adapted from (1)), we take the time derivative of this population,

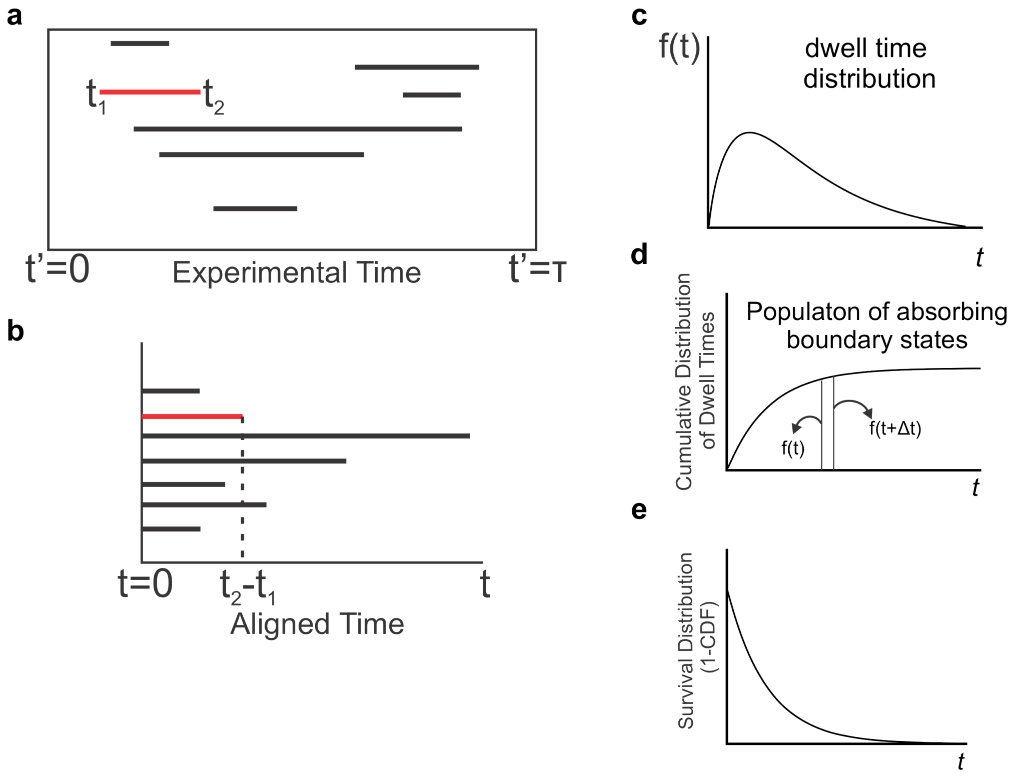


**Supplementary Note Fig. 1: Survival distribution calculation**. (**a**) Experimentally, slow events are seeing as traces in a kymograph. (**b**) These traces are aligned on a new time t and the distribution of lengths corresponds to the dwell time distribution $f(t)$ (**c**) . Shown in red a sample trace before and after time alignment. (**d**) The CDF of $f(t)$ corresponds to the normalized population of an absorbing boundary state. (**e**) (1-CDF) corresponds to the survival distribution $\hat{D}$.

$$f(t) = \frac{d}{dt} \sum_{i \in boundaries} p_i(t) \tag{1}$$

$f(t)$ can be visualized as the probability distribution of experimental track lengths of TFs entering a bound state and evolving independently from a registered time $t = 0$, until they transition to an absorbing boundary state, at which time they leave the bound state (2). The cumulative distribution of $f(t)$ is calculated (Supplementary Note Fig. 1d) and $1 - CDF$ corresponds to the survival distribution ($\hat{D}$, Supplementary Note Fig. 1e) .

#### 1.2 Revised Bi-Exponential Model

We consider an idealized system containing a fixed total number of TFs, each of which can be found in one of three states: Slow (s), Fast (f) and Diffusive (d) (Supplementary Note Fig. 2a) In this model, transitions between s and f states are forbidden. When a transcription factor is either in state s or f, it can be observed experimentally as a trace. When a TF transitions to the state d, its experimental observation will stop and this state plays the role of an absorbing boundary state.

Let $S(t)$ and $F(t)$ denote the number of TFs in the state s and f at the aligned time $t$, respectively. These functions will decrease monotonically to 0 from the initial values $S_0$ and $F_0$, $S_0 + F_0 = N$, which correspond to the total number of TFs in the s and f state during the experiment, respectively. To obtain the survival distribution, the probability that a particle stayed bound (either in state s or f) for a time $t$ or longer, the population of absorbing boundary states needs to be calculated. The latter (population of absorbing boundary states), corresponds to transitions from s to d or from f to d, which denotes the following events:

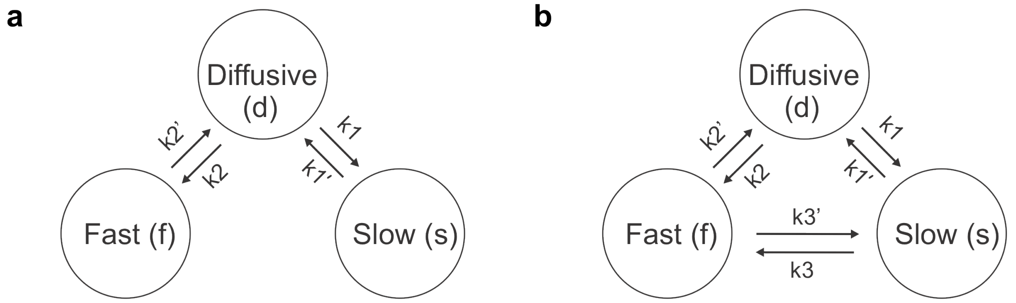


**Supplementary Note Fig. 2: Double Exponential and Kinetic Models**. (**a**) Schematic representation of the double exponential model. Transitions between specific (s) and non-specific binding (f) are forbidden. (**b**) Kinetic model allows for transitions between non-specific and specific binding.

$$s \xrightarrow{k_1'} d \quad f \xrightarrow{k_2'} d \tag{2}$$

Let $D(t)$ denote the number of experimentally observed transcription factors that transitioned from any bound state (s or f) to the diffusive state up to an aligned time $t$, with $D(0) = 0$. This population accumulates over time and the dwell time distribution corresponds to the time derivative of this population. Thus,

$$\frac{dS}{dt} = -k_1'S \quad \frac{dF}{dt} = -k_2'F \tag{3}$$

$$\frac{dD}{dt} = k_1'S + k_2'F \tag{4}$$

Then,

$$S = S_0 e^{-k_1' t}, \quad F = F_0 e^{-k_2' t} \tag{5}$$

$$\frac{dD}{dt} = S_0 k_1' e^{-k_1' t} + F_0 k_2' e^{-k_2' t} \tag{6}$$

where $\frac{1}{N}\frac{dD}{dt}$ corresponds to the dwell time distribution ($f(t)$). The dwell time distribution depends on the initial populations of $S$ and $F$. These correspond to the relative populations entering a cycle in the dwell time ($D \to S$ or $D \to F$), the proportion of traces either in state s or f. Traces in state s or f are born randomly with proportions $\frac{k_1}{k_1+k_2}$ and $\frac{k_2}{k_1+k_2}$ respectively (Supplementary Note Fig 2A).

Then, $S_0 = \frac{Nk_1}{k_1+k_2}$ and $F_0 = \frac{Nk_2}{k_1+k_2}$ which corresponds to the initial values of $S(t)$ and $F(t)$.

Finally, the dwell time distribution is given by (Supplementary Note Fig 1c):

$$\boxed{f(t) = \frac{1}{N}\frac{dD}{dt} = \frac{1}{k_1 + k_2}\left(k_1 k_1' e^{-k_1' t} + k_2 k_2' e^{-k_2' t}\right)} \tag{7}$$

The Survival distribution ($\hat{D} = \int_t^\infty \frac{dD}{du} du$) is given by (Supplementary Note Fig 1e):

$$\boxed{\hat{D} = \frac{1}{k_1 + k_2}\left(k_1 e^{-k_1' t} + k_2 e^{-k_2' t}\right)} \tag{8}$$

$\frac{1}{k_1'}$ and $\frac{1}{k_2'}$ correspond to the expected residence time of the slow and fast component respectively. After calculating the survival distribution, we are interested in finding the proportions of TFs in state s and f in steady state.

At steady state, the slow and fast component populations ($\bar{S}$ and $\bar{F}$) are calculated as follows (Supplementary Note Fig 2a):

$$\frac{d\bar{S}}{dt'} = k_1\bar{D} - k_1'\bar{S} = 0 \rightarrow \bar{S} = \frac{k_1}{k_1'}\bar{D} \tag{9}$$

$$\frac{d\bar{F}}{dt'} = k_2\bar{D} - k_2'\bar{F} = 0 \rightarrow \bar{F} = \frac{k_2}{k_{2'}}\bar{D} \tag{10}$$

The slow and fast component proportions ($\hat{S}$ and $\hat{F}$) are given by:

$$\boxed{\hat{S} = \frac{k_1k_2'}{k_1k_2' + k_1'k_2}; \;\; \hat{F} = \frac{k_1'k_2}{k_1k_2' + k_1'k_2}} \tag{11}$$

Previously (3–7), the survival distribution was phenomenologically fitted to $\hat{D} = \left(f_1e^{-k_1't} + (1-f_1)e^{-k_2't}\right)$ and $f_1$, $(1-f_1)$ were interpreted as the slow and fast component proportions at steady state contrary as the values found in equation 11. The derivation of the survival distribution shows that $f_1 = \frac{k_1}{k_1+k_2}$ represents the proportion of traces in the s state during the experimental observation and not their steady state populations.

#### 1.3 Kinetic Model

We next extended the bi-exponential model so that transitions between the slow and fast components are allowed but indistinguishable. $S, F$ and $D$ defined as in 1.2 (Supplementary Note Fig. 2b). If a TF is in a non-specific bound state (f, diffusing or hopping on the DNA), and it binds to a specific site (s), this transition cannot be observed due to the spatial resolution limit (∼30nm, ∼3 nucleosomes). The particle will appear bound regardless of the number of transitions between the fast and the slow component inside the resolution limited volume. The diffusive state corresponds to an absorbing boundary state (Supplementary Note Fig. 2b), as before. Then, the dwell time distribution $\frac{dD}{dt}$ can be calculated as follows:

$$d \xleftarrow{k_1'} s \underset{k_3'}{\overset{k_3}{\rightleftharpoons}} f \xrightarrow{k_2'} d \tag{12}$$

$$\frac{dS}{dt} = -(k_1' + k_3)S + k_3'F \quad \frac{dF}{dt} = -(k_3' + k_2')F + k_3S \tag{13}$$

$$\frac{dD}{dt} = k_1'S + k_2'F \tag{14}$$

This can be solved in matrix form as:

$$\begin{pmatrix} \frac{dS}{dt} \\ \frac{dF}{dt} \end{pmatrix} = \begin{pmatrix} -(k_1' + k_3) & k_3' \\ k_3 & -(k_3' + k_2') \end{pmatrix} \begin{pmatrix} S \\ F \end{pmatrix} \tag{15}$$

The solution will be given by:

$$\begin{pmatrix} S \\ F \end{pmatrix} = C_1 \begin{pmatrix} \alpha_1 \\ \beta_1 \end{pmatrix} e^{\lambda_1 t} + C_2 \begin{pmatrix} \alpha_2 \\ \beta_2 \end{pmatrix} e^{\lambda_2 t} \tag{16}$$

Where $\lambda_i$ are the eigenvalues and $\begin{pmatrix} \alpha_i \\ \beta_i \end{pmatrix}$ the corresponding eigenvectors of the matrix in equation 15. $C_1$ and $C_2$ are calculated from the initial populations of the state s and f. These populations are given by $\frac{Nk_2}{k_1+k_2}$ and $\frac{Nk_1}{k_1+k_2}$ for states f and s respectively which corresponds to the number of TFs that entered a bound state through state s or f respectively. Then, the dwell time distribution is given by:

$$\boxed{f(t) = k_2'F(t) + k_1'S(t)}. \tag{17}$$

The survival distribution is calculated as:

$$\hat{D} = \int_t^\infty f(u)du$$

with $S, F$ as defined in equation 16. The survival distribution corresponds then to a sum of two exponentials similar to the double exponential model but with exponential parameters ($\lambda_1$ and $\lambda_2$) that depend on the rate constants of the process. Therefore, double exponential fits to the experimental data cannot be directly used to extract the kinetic rates of the underlying process.

#### 1.4 Diffusion and binding on a chain

A number of theoretical studies have posited that the process of TF binding to its cognate site on DNA involves a combination of bulk diffusion in the nucleus, 1-d sliding along the DNA, hopping and translocation, and have derived the search time subject to various conditions. In this extension of our basic model and to account for a multiplicity of fast bound states, we can model TF searching for a specific site on the DNA by assuming the DNA to be a circular chain composed of specific sites and non-specific sites (Supplementary Note Fig. 3a). The assumption in the following derivation is that the number of non-specific sites on the DNA is much larger than specific sites. This is a biologically reasonable assumption as only a few tens of thousands specific sites are bound by any TF according to genome wide studies (8, 9), while the entire genome contains millions of "other" potential chromatin binding sites. A TF binds stochastically to any site on the DNA and diffuses around the chain with a certain probability of unbinding from any state. The hopping rate from a non-specific site to another site (specific or non-specific), is given by $k_1$, the rate of hopping from a specific site to a non-specific or dissociation to the bulk is given by $k_2$ and the rate of dissociation from a non-specific site into the bulk is given by $k_3$.

Thus the empirically observed transitions of TF dissociations can result from two possibilities: (i) a TF begins at a random location in the chain and diffuses along the DNA on non-specific sites and unbinds to the bulk diffusive state (d) before finding the specific site; (ii) a TF diffuses along the DNA until reaches a specific site, and it unbinds from the DNA from either a specific site or a non-specific site.

##### 1.4.1 A TF does not find a target

The case when a TF does not find a target is dynamically equivalent to a TF binding to a non-specific site with the possibility of diffusing to yet another non-specific site (Supplementary Note Fig. 3b). Due to the self-similarity of the chain, all non-specific sites are indistinguishable. Here, we assume that it is equally probable for a TF to jump between non-specific sites and unbind from a non-specific site, i.e $k_1 = k_3$. Applying the absorbing boundary method, the dwell time distribution can be easily found:

$$\frac{dD}{dt} = k_1 (P_1 + P_2) \tag{18}$$

where $P_1(t)$ and $P_2(t)$ correspond to the number of traces at time $t$ that started in state 1 or 2 respectively.

$$\frac{dP_1}{dt} = -2k_1P_1 + k_1P_2, \ \frac{dP_2}{dt} = -2k_1P_2 + k_1P_1 \tag{19}$$

$$\rightarrow \begin{pmatrix} \frac{dP_1}{dt} \\ \frac{dP_2}{dt} \end{pmatrix} = k_1 \begin{pmatrix} -2 & 1 \\ 1 & -2 \end{pmatrix} \begin{pmatrix} P_1 \\ P_2 \end{pmatrix} \tag{20}$$

$$\rightarrow \begin{pmatrix} P_1 \\ P_2 \end{pmatrix} = C_1 \begin{pmatrix} 1 \\ 1 \end{pmatrix} e^{-k_1 t} + C_2 \begin{pmatrix} -1 \\ 1 \end{pmatrix} e^{-3k_1 t} \tag{21}$$

After applying initial conditions for $P_1$ and $P_2$ (each non specific site is equally probable, $C_1 = \frac{1}{2}$ and $C_2 = 0$), we find that the normalized dwell time distribution is given by:

$$\boxed{\frac{d\tilde{D}}{dt} = k_1 e^{-k_1 t}}, \tag{22}$$

as expected for a Poisson process. The exponential term due to the dissociation of the TF from non-specific sites along the DNA dominates the internal diffusion of the TF along the chain.

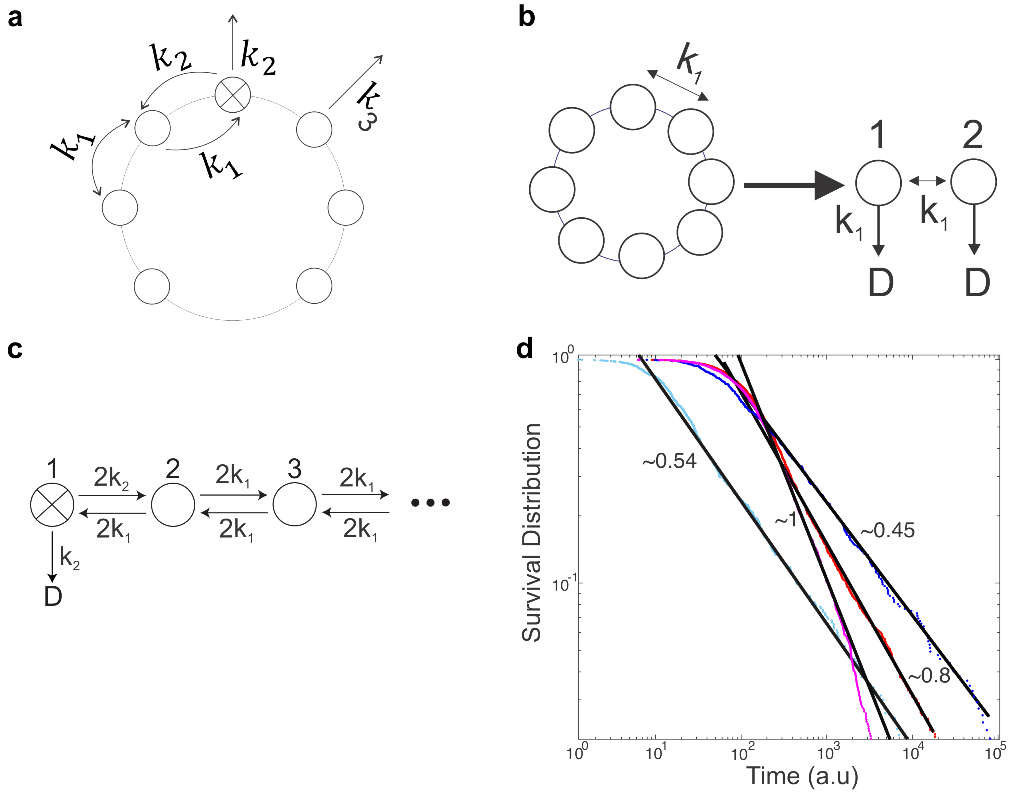


**Supplementary Note Fig. 3: Diffusion and binding along a chain leads to power law behaviour of dwell times**. (**a**) Schematic of TF diffusion on DNA represented as a circular chain that contains either non-specific sites (empty circles) or specific sites (crossed circle). (**b**) Equivalent state diagram for the condition when a TF does not find a specific site. The rates for diffusing along the chain and dissociating from the chain are set to $k_1$. (**c**) Equivalent state diagram for the case when the TF finds a specific site and unbinds from this site to the bulk. (**d**) Survival distribution of TF calculated using stochastic simulations. TFs bind uniformly around 20 sites away from the specific site. The simulations show different power law decays depending on different choice of parameters. For power-law of 0.45, $k_1 = 1$, $k_2 = 0.1$ and $k3 = 1$; for power-law of 0.54, $k_1 = 10$, $k_2 = 0.1$ and $k3 = 1$; for power-law of 0.8, $k_1 = 10$ $k_2 = 0.01$ and $k_3 = 0.01$; for power-law of 1, $k_1 = 10$ $k_2 = 0.01$ and $k_3 = 0.0001$.; for power-law 1.5, $k_1 = 1$ $k_2 = 0.01$ and $k_3 = 0.001$

##### 1.4.2 A TF does find its target

Let $P_n(t)$ denote the probability of being in state $n$ at time $t$. We will consider (without loss of generality) that $k_1 = 0.5$ and $\frac{k_2}{k_1} = \alpha$; $k_1$ will impose the time scale of the processes shown in Supplementary Note Fig. 3c. We are considering an infinite linear chain since a TF factor after finding its specific site will either move to a non-specific site to the right or left with equal probability and since we are considering a long chain, the right-side chain will be the reflection of the left-side chain. To solve this case analytically we are considering the special case where unbinding is only possible from a specific site. The master equation associated with this process is given by:

$$\frac{\partial P_1(t)}{\partial t} = P_2(t) - 1.5\alpha P_1(t) \quad \text{for } n = 1 \tag{23}$$

$$\frac{\partial P_2(t)}{\partial t} = \alpha P_1(t) + P_3(t) - 2P_2(t) \quad \text{for } n = 2 \tag{24}$$

$$\frac{\partial P_n(t)}{\partial t} = P_{n-1}(t) + P_{n+1}(t) - 2P_n(t) \quad \text{for } n > 2 \tag{25}$$

Site 1 corresponds to the specific site and the initial condition is such that $P_n(0) = \delta_{n,1}$.

Using the property of the Modified Bessel functions of first kind, $\frac{\partial I_n(t)}{\partial t} = \frac{1}{2}\left(I_{n-1}(t) + I_{n+1}(t)\right)$. It is easy to see that $P_n(t) = e^{-2t} I_{n+m}(2t)$ are solutions for 25, with $m \in \mathbb{N}$. Let's consider the linear combination of these particular solutions:

$$\boxed{P_n(t) = \sum_{m=0}^{\infty} A_m^n e^{-2t} I_m(2t)} \tag{26}$$

Applying the general differential equation for $n > 2$ (25), we get the following conditions on the coefficients:

$$\boxed{A_2^n + 2A_0^n = A_1^{n+1} + A_1^{n-1}} \tag{27}$$

$$\boxed{A_{m+1}^n + A_{m-1}^n = A_m^{n-1} + A_m^{n+1}} \quad m \neq 1 \tag{28}$$

Without loss of generality, we can set $A_1^4 = 0$. This implies that:

$$\boxed{A_{m+1}^n = A_m^{n-1};\ \ A_{m-1}^n = A_m^{n+1}} \tag{29}$$

Applying the initial condition that $P_n(0) = \delta_{n1}$, the solution is given by:

$$\boxed{P_n(t) = e^{-2t}\left\{A_{n-1}^n I_{n-1}(2t) + \sum_{m \neq n-1} A_m^n I_m(2t)\right\}} \tag{30}$$

with $A_0^1 = 1,\ A_0^n = 0,\ n > 1$. Using this solution as ansatz for 24:

$$P_1(t) = e^{-2t}\left\{I_0(2t) + \sum_{m=1} A_m^1 I_m(2t)\right\} \tag{31}$$

$$P_2(t) = e^{-2t}\left\{A_1^2 I_1(2t) + \sum_{m=1} A_{m+1}^2 I_{m+1}(2t)\right\} \tag{32}$$

$$\frac{\partial P_2(t)}{\partial t} = \alpha P_1(t) + P_3(t) - 2P_2(t) \tag{33}$$

for simplicity the notation $I_m(2t) \equiv I_m$ is used from now on. Then,

$$I_0\left(A_1^2-\alpha\right)-I_1\left(\alpha A_1^1+A_1^3-A_2^2\right) \tag{34}$$

$$=\sum_{m=3} I_m\left(A_m^3+\alpha A_m^1-A_{m-1}^2-A_{m+1}^2\right)+I_2\left(A_2^3+\alpha A_2^1-A_1^2-A_3^2\right)$$

This implies:

$$\boxed{A_1^2=\alpha;\quad A_1^3=A_2^2-\alpha A_1^1} \tag{35}$$

Applying $A_{m+1}^n=A_m^{n-1}$:

$$A_3^2=\alpha A_2^1 \tag{36}$$

$$\boxed{A_{m+1}^2=\alpha A_m^1\quad m>2} \tag{37}$$

Applying the ansatz to 24, we find:

$$(1.5\alpha-2+A_1^1)I_0+(2-A_1^2)I_1=\sum_{m=1} I_m\left(A_m^2-1.5\alpha A_m^1-A_{m+1}^1-A_{m-1}^1+2A_m^1\right) \tag{38}$$

applying $A_{m+1}^2=\alpha A_m^1$ , $A_1^2=\alpha$, $A_2^2=A_1^3+\alpha A_1^1$, $A_3^2=\alpha A_2^1$ and setting without loss of generality $A_1^3=0$,

$$\Rightarrow (1.5\alpha-2+A_1^1)I_0+I_1(2-\alpha-A_1^1(2-1.5\alpha)+A_2^1)=\sum_{m=4} I_m\left((2-1.5\alpha)A_m^1-A_{m+1}^1+(\alpha-1)A_{m-1}^1\right) \tag{39}$$

$$+I_2\left(A_2^2-1.5\alpha A_2^1-A_3^1-A_1^1+2A_2^1\right)+I_3\left(A_3^2-1.5\alpha A_3^1-A_4^1-A_2^1+2A_3^1\right)$$

Then,

$$\boxed{A_4^1=(\alpha-1)A_2^1+(2-1.5\alpha)A_3^1} \tag{40}$$

$$\boxed{A_3^1=A_1^3+A_1^1(\alpha-1)+A_2^1(2-1.5\alpha)} \tag{41}$$

$$\boxed{A_1^1=2-1.5\alpha;\ A_2^1=\alpha-2+A_1^1(2-1.5\alpha);\ A_{m+2}^1=(2-1.5\alpha)A_{m+1}^1+(\alpha-1)A_m^1} \tag{42}$$

Finally,

$$\boxed{P_1(t)=e^{-2t}\left\{I_0(2t)+(2-1.5\alpha)I_1(2t)+\left[\alpha-2+A_1^1(2-1.5\alpha)\right]I_2(2t)+\sum_{m=1}^{\infty}A_{m+2}^1 I_{m+2}(2t)\right\}} \tag{43}$$

In the limit when $t\gg 1$, we can apply the following approximation (10):

$$I_n(2t)\approx(4\pi t)^{-1/2}e^{2t}\left[1-\frac{(4n^2-1)}{16t}+\frac{\left(4n^2-1\right)\left(4n^2-9\right)}{2!\,(16t)^2}+...\right] \tag{44}$$

We want to derive the asymptotic behaviour of the solution for $P_1(t)$. For this, we need to calculate the leading terms of the expansion for $I_n(2t)$. First, we are going to calculate the behavior of the leading order of $P_1(t)$, terms of the form $(4\pi t)^{-1/2}$. Assuming that $\alpha\neq 1$:

$$P_1(t)\approx(4\pi t)^{-1/2}\sum_{m=0}^{\infty}A_m+\mathcal{O}(t^{-3/2})$$

$$\sum_{m=0}^{\infty} A_m = A_0 + A_1 + A_2 + A_3 + A_4 + \sum_{m=3}^{\infty} A_{m+2} \tag{45}$$

Using 42,

$$\sum_{m=0}^{\infty} A_m = A_0 + A_1 + A_2 + A_3 + A_4 + \sum_{m=4}^{\infty}(2 - 1.5\alpha)A_m^1 + \sum_{m=3}^{\infty}(\alpha - 1)A_m^1 \tag{46}$$

$$\Rightarrow 0.5\alpha \sum_{m=0}^{\infty} A_m = 0.5\alpha(A_0 + A_1 + A_2) + A_3(1.5\alpha - 1) + A_4 \tag{47}$$

Replacing the values for $A_0$, $A_1$, $A_2$, $A_3$ and $A_4$; it is easy to see that:

$$\Rightarrow \boxed{\sum_{m=0}^{\infty} A_m = 0} \tag{48}$$

Which implies that terms of order $t^{-1/2}$ vanish for the solution of $P_1(t)$.We need to calculate the behaviour of the second leading term of the form $\frac{(4\pi t)^{-1/2}}{16t}$ (terms of the order $t^{-3/2}$) which will dominate the asymptotic behaviour of $P_1(t)$. It is easy to see in a similar way that for $\sum_{m=0}^{\infty} A_m$, that

$$\boxed{0 < \sum_{m=0}^{\infty} A_m \frac{(4m^2 - 1)}{16} < \infty} \tag{49}$$

This implies that the asymptotic behavior of $P_1(t)$ will be dominated by a term of the form $\frac{(4\pi t)^{-1/2}}{16t}$:

$$\boxed{P_1(t) \propto t^{-3/2}} \tag{50}$$

Finally, the dwell time distribution $\tilde{D}$ is given by:

$$\frac{d\tilde{D}}{dt} = 0.5\alpha P_1(t) \propto 0.5\alpha t^{-3/2} \tag{51}$$

The survival distribution $\hat{D}$ in the limit $t \gg 1$:

$$\boxed{\hat{D}(t) \propto 0.5\alpha t^{-1/2}}, \tag{52}$$

The survival distribution for such a chain will be the sum of an exponential term due to non-specific binding with a fast dwell time and a term of the form $t^{-1/2}$ as described above ($t \gg 1$). The asymptotic behavior for longer times compared to diffusion along the DNA (random walk along non-specific sites) will therefore show power-law behavior. This suggests that the phenomenological fits of survival time distributions by a mixture of exponentials may have to be modified depending on TF affinity and dynamics.

This analytical result can be applied to transcription factors that undergo multiple conformational changes or interact with multiple protein complexes before unbinding from a specific site. Such a process will produce asymptotically power-law distributed dwell times for specific binding.

Conceptually, our calculation shows that diffusion on the DNA as illustrated in Section 1.4 produces asymptotic power-law behavior for the survival distribution of TFs. This diffusion on the DNA may be seen as a broad distribution of effective affinities depending on how many non-specific targets the TF visited prior to binding to a specific site with each specific binding event having a distinct effective binding affinity.

The next section will explore computationally a more general process where unbinding is allowed from non-specific sites.

##### 1.4.3 Simulating a Complete Model

We used the Gillespie algorithm (11) to simulate the residence time of a TF binding to a specific target in the background of multiple non-specific sites.We modeled the chromatin environment as a circular chain of 2000 sites, with 1999 non-specific and a single specific site (Supplementary Note Fig. 3). TFs were initially allowed to bind at a randomly chosen non-specific site at most 20 sites away from the specific site, since TFs bound farther away contribute negligibly to the final residence time distribution due to the low probability of finding the specific site before unbinding. TFs were allowed to diffuse to a neighboring site with the rate constants as shown in Supplementary Note Fig. 3a or dissociate from the specific site at a rate $k_2$ or from a nonspecific site at a rate $k_3$. The dwell time for an iteration corresponds to the time interval between the initial condition of binding at a random site and unbinding of the TF. The simulation was terminated when 1000 specific binding events occurred (multiple specific binding for the same TF in a single iteration were counted as a single binding event) or after $1 \times 10^7$ time points (a.u.). The survival distribution was calculated using the Kaplan-Meier estimate for the empirical cumulative distribution function. We find that the survival time distribution clearly shows a power law dependence for different combinations of $k_1$, $k_2$ and $k_3$ when $k_2 < k_1$ (Supplementary Note Fig. 3d). Unbinding from a non-specific site was allowed after a TF binds to a specific site. The power-law exponent is largely determined by the search time or the effective diffusion on the DNA, before finding the specific site based on exploration of different parameters through simulations. Conceptually, our simulation implies that since a TF can visit multiple non-specific sites before binding to the specific site and subsequently unbinding, the effective survival distribution resembles the situation in which the TF encounters sites with a broad distribution of affinities. This suggests that the phenomenological fits of survival time distributions by a mixture of exponentials may have to be modified depending on TF affinity and dynamics.

#### 1.5 Broad distribution of Binding Affinities

Independent of diffusion along the DNA, the affinity landscape in the nucleus may be highly heterogeneous resulting in a broad distribution of binding affinities contrary to the assumption that TF dynamics on chromatin results from well-separated and narrow distributions of specific and non-specific binding (Fig. 5). Given the heterogeneity in local organization and nuclear structure, TF binding sites on chromatin can be viewed as a collection of traps with a distribution of trap depths (analogous to binding affinities, Figure 4H). Each trap can be viewed as a potential energy well of depth $\Delta E$, which is related to the affinity as $k(T) = k_0 e^{-\Delta E/k_B T}$, where $k_0$ is the bare transition rate (Arrhenius equation). The dwell time distribution of a TF in a particular well of depth $\Delta E$ is given by $f_{\Delta E}(t) = k(T)e^{-k(T)t}$. If the energy landscape across the nucleus is described by a distribution $P(\Delta E)$, then the overall dwell time distribution observed in an experiment across all different sites ($f(t)$) will take the form:

$$\boxed{f(t) = \int k_0 e^{-\Delta E/k_B T} e^{-k_0 e^{-\Delta E/k_B T} t} P(\Delta E) d\Delta E} \tag{53}$$

Under very general conditions, Bouchaud et al. have shown that for such finite disordered systems, there exist a family of energy distributions $P(\Delta E)$, for which the distribution of dwell times asymptotically approaches a power law (12, 13). As an example, the well-known Random Energy model and the Sherrington-Kirkpatrick model of spin glasses from physics have energy landscape in which the distribution of deep traps (large $\Delta E$) is given by:

$$P(\Delta E) = \frac{f_0}{k_B T} e^{-x\Delta E/k_B T} \tag{54}$$

where $x$ is a temperature dependent parameter between 0 to 1. For the case of TF binding, this parameter $x$ can be seen as the interaction strength of different types of TFs to a specific site described by a well with energy depth $\Delta E$. Using this distribution for the energy landscape, the dwell time distribution can be calculated:

$$f(t) = \frac{f_0 k_0}{k_B T} \int_0^\infty e^{-\Delta E/k_B T(1+x)} e^{-k_0 e^{-\Delta E/k_B T} t} d\Delta E \tag{55}$$

let's $u = e^{-\Delta E/k_B T}$ and $t' = k_0 t$:

$$f(t) = f_0 k_0 \int_0^{t'} u^x e^{-e^u t'} du \tag{56}$$

setting $ut' = v$,

$$f(t) = f_0 k_0 \int_0^{t'} \left(\frac{v}{t'}\right)^x e^{-v} \frac{dv}{t'} \tag{57}$$

$$\Rightarrow f(t) = \frac{f_0 k_0}{t'^{(1+x)}} \int_0^{t'} v^x e^{-v} dv \tag{58}$$

Using the definition for the incomplete gamma function ($\Gamma(a, x) \equiv \int_x^\infty t^{a-1} e^{-t} dt$)and using $t' = k_0 t$

$$f(t) = t^{-(1+x)} k_0^{-x} f_0 \left[\Gamma(x+1, k_0 t) - \Gamma(x+1, 0)\right] \tag{59}$$

We are interested in the asymptotic behavior of $f(t)$. For $t \gg 1$:

$$f(t) \approx -t^{-(1+x)} k_0^{-x} f_0 \Gamma(x+1, 0) \tag{60}$$

Let $C(x) = k_0^{-x} f_0 \Gamma(x+1, 0)$ which is a time independent quantity ($C(x) < 0$),

$$f(t) \approx -C(x) t^{-(1+x)} \tag{61}$$

Finally, the asymptotic behavior of the survival distribution $\hat{D}(t)$ ($t \gg 1$:) is given by:

$$\boxed{\hat{D}(t) \propto t^{-x}} \tag{62}$$

Thus, the survival distribution exhibits asymptotic power-law behavior over a range of parameters that are experimentally plausible.

##### 1.5.1 Simulation

To simulate a broad distribution of binding affinities, the chromatin was modeled as a chain of 10000 sites. The depth of potential well for each site was randomly chosen from an exponential distribution with a mean of $\frac{x}{k_B T}$ (x was set up to 0.5). The affinity distribution for the site then becomes $k = k_0 e^{-\Delta E/k_B T}$ and the resulting dwell time distribution for any particular well was randomly generated from an exponential distribution with mean $1/k$. The Survival distribution was then calculated using the Kaplan-Meier estimator.
